# Impact of Common Modifications on the Antigenic Profile and Glycosylation of Membrane-Expressed HIV-1 Envelope Glycoprotein

**DOI:** 10.1101/2023.06.02.543368

**Authors:** Tommy Tong, Alessio D’Addabbo, Jiamin Xu, Himanshi Chalwa, Albert Nguyen, Paola Ochoa, Max Crispin, James M Binley

## Abstract

Recent HIV-1 vaccine development has centered on “near native” soluble envelope glycoprotein (Env) trimers. These trimers are artificially stabilized laterally (between protomers) and apically (between gp120 and gp41). These same stabilizing mutations have been leveraged for use in membrane-expressed Env mRNA vaccines, although their precise effects in this context are unclear. To address this question, we investigated the effects of Env mutations expressed on virus-like particle (VLP) in 293T cells. Uncleaved (UNC) trimers were laterally unstable upon gentle lysis from membranes. However, gp120/gp41 processing improved lateral stability. Due to inefficient gp120/gp41 processing, UNC is incorporated into VLPs. A linker between gp120 and gp41 (NFL) neither improved trimer stability nor its antigenic profile. An artificially introduced enterokinase cleavage site allowed processing post-expression, resulting in increased trimer stability. Gp41 N-helix mutations I559P and NT1-5 both imparted lateral trimer stability, but concomitantly reduced gp120/gp41 processing and/or impacted V2 apex and interface NAb binding. I559P consistently reduced recognition by HIV+ donor plasmas, further supporting antigenic differences. Mutations in the gp120 bridging sheet failed to stabilize membrane trimers in a pre-fusion conformation, reduced gp120/gp41 processing and exposed non-neutralizing epitopes. Reduced glycan maturation and increased sequon skipping were common effect of mutations. In some cases, this may be due to increased rigidity which limits access to glycan processing enzymes. In contrast, viral gp120 did not show glycan skipping. We observed a minor species of high mannose glycan only gp160 in particle preparations. This was unaffected by any mutations and instead bypasses normal folding and glycan maturation processes. Including the full gp41 cytoplasmic tail led to markedly reduced gp120/gp41 processing and increased the proportion of high mannose gp160. Remarkably, NAbs were unable to bind to full-length Env trimers. Overall, our findings suggest caution in leveraging mutations to ensure they impart valuable membrane trimer phenotypes for vaccine use.

**AUTHOR SUMMARY:** A vaccine that induces virus-fighting antibodies to block HIV-1 infection remains elusive. To ablate HIV-1 infection, antibodies must bind to authentic envelope (Env) glycoprotein on the virus surface. However, Env can exist in various forms, many of which are relatively easy targets for non-effective antibodies. Therefore, a key challenge of vaccine design is to create pure authentic Env that is unfettered by these other forms of Env. Vaccine research to date has focused largely on stabilizing soluble Env trimers, as this format simplifies purification and translational studies. However, incomplete Env authenticity may blunt the efficacy of this approach. By comparison, the manufacture of particle-based vaccines that express Env *in situ* on membranes is cumbersome. In an alternative approach, lipid particles can deliver mRNA vaccines encoding membrane trimers, bypassing the manufacturing challenges of previous methods. Stabilizing mutations derived from soluble trimers are now being leveraged for membrane trimers. Here, we evaluated the effects of these mutations. Our results show that some mutations alter Env conformation, and therefore might best be omitted from membrane Env vaccines.

## INTRODUCTION

HIV-1 envelope glycoprotein (Env) gp120/gp41 trimers are the natural target of broadly neutralizing antibodies (bNAbs). These trimers use sophisticated strategies to evade bNAbs, including high sequence variation of exposed surfaces, a dense glycan shield, and conformationally dynamic epitopes [1–4]. Env trimers can assume various conformations, ranging from fully open to the closed, pre-fusion state. Many bNAbs exhibit a preference for the pre-fusion state (termed “state 1”) [1,3], supporting efforts to present this form of Env in vaccines.

Mutational studies aimed at stabilizing soluble gp140 laid the groundwork for the first crystal structure of a “near native” trimer [5]. Gp140 was first stabilized apically via an “SOS” disulfide bond between gp120 and gp41 (A501C+T605C) [6], then laterally via a proline mutation in the N helix of gp41 (I559P), collectively termed “SOSIP” gp140 [7]. I559P destabilizes the low energy post-fusion “6 helix bundle” conformation of transmembrane domains, so that the C-helices can self-associate. Alternatively, a linker also situated in the gp41 N-helix (AA548-568, termed UFO) lends lateral stability [8]. To address the challenge of incomplete cellular processing into gp120/gp41, one solution has been to introduce a linker, e.g., NFL, to provide gp120 and gp41 with flexibility to associate in a native-like manner [9,10].

Soluble trimer structures have revitalized vaccine research by providing a template for rational, mutation-based vaccine designs. For example, A433P and DS (I201C+A433C to cross-link the β3 and β21 strands of gp120) both prevent soluble trimers from adopting a CD4-induced state [11]. Other mutation strategies include: i) leveraging residues from strains that form well-folded gp140 trimers [12–20], ii) rendering trimers vulnerable to bNAb precursors [21,22], and iii) stabilizing the pre-fusion state by improving electrostatic interactions [23].

Vaccines that present authentic trimers in their natural, membrane context might be our best hope to elicit bNAbs. Accordingly, we have been developing virus-like particles (VLPs) expressing trimers as a vaccine platform [24–26]. Native trimers are generated from an uncleaved (UNC) gp160 precursor. The conformational flexibility of UNC gp160 is essential for it to reach its mature form, facilitating glycan addition/maturation, disulfide isomerization until canonical disulfides form, followed by signal peptide removal and gp120/gp41 processing [27,28]. It is therefore not surprising that UNC exists in a variety of oligomeric states that are sensitive to non-NAbs [1,29–34].

To date, various membrane trimer mutants (SOS, I559P and UNC), like their soluble gp140 counterparts, mostly adopt relatively “open” trimer states 2 and 3, recognized by non-NAbs [3,35–42]. On the other hand, native membrane gp120/gp41 trimers, but not UNC, are exclusively bound by NAbs [30,33,34]. Incomplete gp120/gp41 processing is a consistent unwanted feature of membrane Env expressed on VLPs, pseudovirions and even infectious molecular clones expressed in 293T cells [30,43–45]. Non-functional Env comes in a variety of forms. While some UNC gp160 acquires complex glycans, a fraction acquires only high mannose glycans, apparently bypassing normal glycosylation [32,34,39,46]. Furthermore, gp120 shedding leaves behind gp41 stumps [30].

mRNA vaccines have great potential to simplify the otherwise challenging translation of membrane-expressed trimer vaccine concepts into the clinic [47–50]. So far, data suggests that mRNA-expressing membrane trimers outperform soluble trimers [10,51–55]. Mutations that were previously used for soluble trimer vaccines have been leveraged in corresponding membrane trimers [51-53,56-58]. A phase I trial of 3 mRNA-based HIV vaccines is now underway (HVTN302), 2 of which express membrane Env and carry various mutations including SOS, I559P, NFL-like linker, cytoplasmic tail truncation, and mutations to improve antigenicity and thermal stability [59]. However, the effects of these stabilizing mutations may differ in the context of membrane trimers. While some mutations, for example SOS, are widely adaptable, others may have context-dependent effects [22]. The DS mutation may decrease gp120/gp41 processing of membrane trimers [36]. Similarly, the effects of I559P and UFO might differ for membrane trimers if they are already stable in membrane context [10,30,38,53,60]. These challenges are compounded by the co-expression of non-functional Env isoforms [30]. This is important because mRNA expression cannot “filter out” this non-functional Env that might dampen antibody responses away from the intended native trimer targets.

Previously, a panel of HIV Env mutants including SOS, I559P, A433P, DS, NFL, UFO, both alone and in combination, were evaluated antigenically using MAbs to down select lead constructs for mRNA delivery [10]. However, it remains unclear how these mutations impact stability, gp120/gp41 processing, expression, glycosylation and non-functional Env. To fill this knowledge gap and to identify valuable mutation(s) for future mRNA vaccine studies, we compared various mutants alone and in combination in the context of VLP-expressed membrane trimers. Our data suggest that modifications variably impacted Env expression, gp120/gp41 processing, glycosylation, antigenicity, and stability in the lateral and apical planes. These findings can assist in making informed modifications in membrane-expressed vaccine constructs.

## RESULTS

### Expression and maturation of membrane trimer mutants

We typically make VLPs expressing SOS mutant Env because it eliminates gp120 shedding. In addition, we truncate the gp41 cytoplasmic tail (gp160ΔCT) to improve expression and gp120/gp41 processing [30]. The resulting mutant is measurably infectious and typically exhibits a tier 2 profile [30,61]. Other mutations have been developed to stabilize soluble gp140 trimers in a “closed” conformation resembling native spikes. These include deletions, linkers, engineered disulfides and proline substitutions. It is unclear which of these modifications are needed for optimal NAb induction by mRNA vaccine-encoded Env *membrane* trimers [10,52–54]. To address this question, we generated a panel of mutants, including SOS, NFL, I559P, UFO, A433P, DS, A328G [25] and enterokinase cleavage site (EK) mutants [62], alone and in combination in the JR-FL E168K+N189A gp160ΔCT parent background. These Env mutants, depicted schematically in Fig. 1, were co-expressed with Rev and MuLV Gag plasmids in 293T cells to make VLPs.

**Figure 1.**
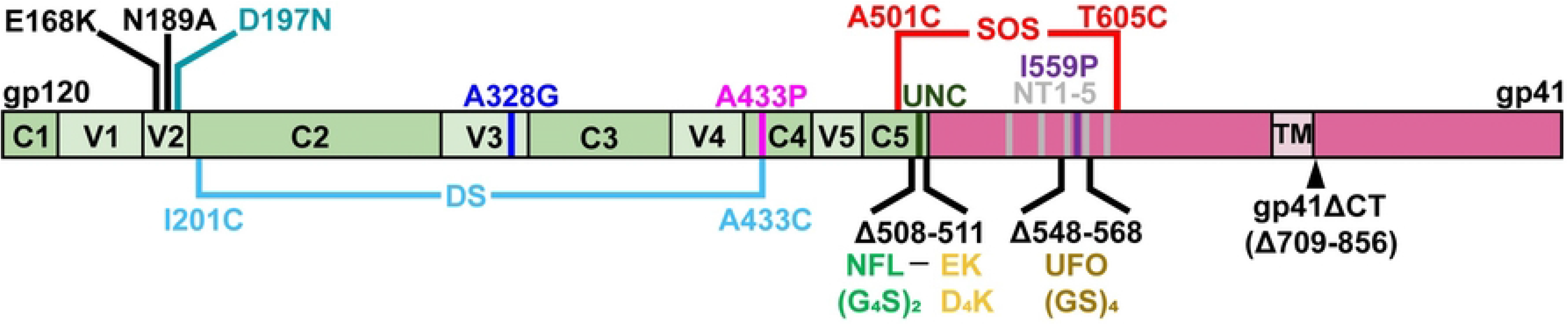
HIV-1 gp160 mutants. A schematic of gp160 showing gp120 (green) variable (V1-V5) domains, constant domains (C1-C5), and gp41 (salmon), including the transmembrane domain (TM). Mutants are shown in different colors that we also use consistently hereafter for clarity. Amino acids are numbered according to HxB2 subtype B reference strain. Gp41 was truncated from position 709 onwards, leaving three amino acids after the TM domain in gp160ΔCT. Most of our JR-FL mutants are based on gp160ΔCT SOS E168K+N189A, unless otherwise stated. NFL is “native flexibly linked” with Gly-Gly-Gly-Gly-Ser sequence repeated twice. EK is an “enterokinase” digestion site with sequence Asp-Asp-Asp-Asp-Lys. UFO is “uncleaved prefusion-optimized” consists of the sequence Gly-Ser repeated four times replacing residues 548-568. The UNC consists of K510S+R511S. The NT1-5 mutant consists of M535I+L543Q+N553S+Q567K+G588R.

A gp120-gp41 linker may grant the flexibility to associate in a “near native” conformation. Thus, linkers could be a partial remedy for incomplete gp120/gp41 processing to improve the consistency of Env products. Accordingly, we included NFL linkers [9] in many of our combination mutants. VLP lysates were first compared in reducing SDS-PAGE-Western blots (Fig. 2). As expected, NFL eliminated gp120 and gp41 bands (missing red and green dots, respectively in lanes 2-10). Consistent with previous studies [30,34,39,43–46,63,64], we detected two forms of gp160: a “mature” gp160 (gp160m, magenta dots) bearing complex glycans and an “immature” gp160 (gp160i, yellow dots) bearing high mannose glycans. In NFL mutants, the gp120 band of the SOS parent (Fig. 2A, lane 1 red dot) was replaced by gp160m (Fig. 2A, lane 2, magenta dot), indicating that, in the absence of a NFL linker, gp160m is partially processed into gp120 and gp41. In contrast, gp160i is unaffected by NFL.

**Figure 2.**
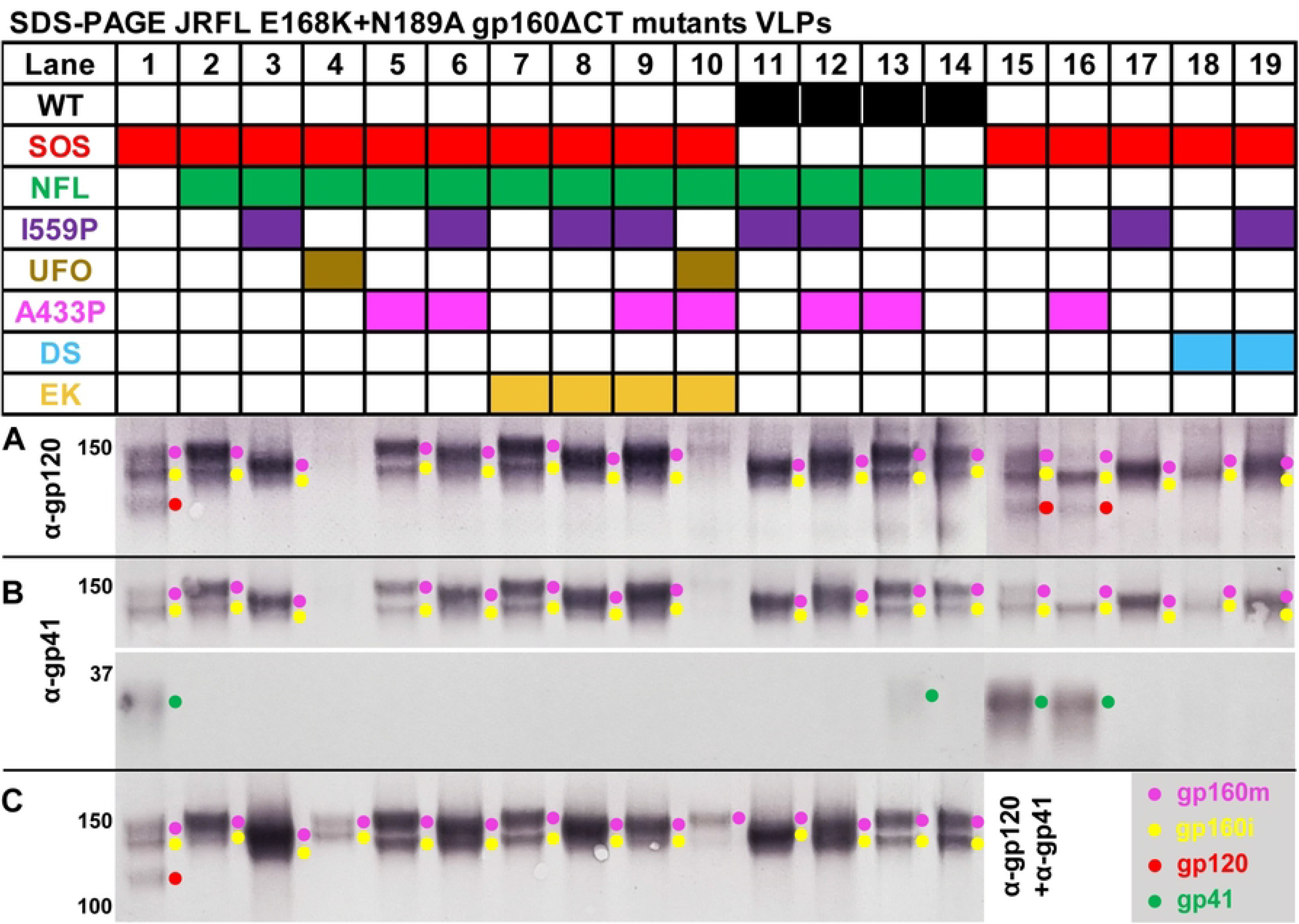
SDS-PAGE-Western blot of gp160ΔCT mutants. VLPs expressing various JR-FL gp160ΔCT E168K+N189A mutant combinations were analyzed by reducing SDS-PAGE-Western blot. Duplicate blots were probed with A) anti-gp120, B) anti-gp41 or C) anti-gp120+gp41 MAb cocktails. Env species are indicated by colored dots.

I559P increased gp160m mobility so that it almost co-migrated with gp160i (Fig. 2, compare lane 2 to lanes 3, 6, 8, 9, 11, 12, 17, and 19). In mutants lacking NFL, I559P impaired gp120/gp41 processing, revealed by the absence of gp120 and gp41 bands (Fig. 2A and B, compare lane 15 to lanes 17 and 19). I559P also improved Env expression (Fig. 2C, compare mutant pairs in lanes 2 and 3, 5 and 6, 7 and 8, 11 and 14, 12 and 13).

Surprisingly, UFO clones expressed very poorly (Fig. 2, lanes 4 and 10), perhaps because they were made in context with NFL. Evidently, these two proximal linkers adversely affect folding. Weak bands were only detected when blots were probed with the full gp120 and gp41 Mab cocktails (Fig. 2C, lanes 4 and 10). The A433P mutant is intended to disrupt the formation of the bridging sheet required for coreceptor binding, thereby stabilizing the trimer in ground state. Here, A433P slowed Env mobility. This was particularly clear when it was used in combination with I559P that increases Env mobility (Fig. 2, compare lanes 3 and 6, 8 and 9, 11 and 12). In the absence of NFL, A433P reduced gp160m, suggesting that it is processed more efficiently (Fig. 2, compare lanes 15 and 16). In contrast, DS impaired gp160 processing and reduced expression (Fig. 2, compare lanes 15 and 18, 17 and 19). High expression was restored when DS and I559P mutants were used together (Fig. 2, lane 19).

The behavior of WT mutants mirrored those of the SOS mutants above (Fig. 2, lanes 11-14). WT NFL exhibited gp160m and gp160i bands (Fig. 2, compare lanes 2 and 14). I559P and A433P mutants increased or decreased gp160m mobility, respectively (Fig. 2, compare lanes 3 and 11, and lanes 5 and 13). I559P and A433P combined led to an intermediate phenotype (Fig. 2, compare lanes 3, 5 and 6, 11-13). A faint gp41 band was detected for the A433P mutant (Fig. 2B, lane 13) suggesting that the NFL is slightly sensitive to host proteases in WT mutants, but not in SOS and I559P mutants. Finally, NFL mutants containing an enterokinase site (EK+) were comparable to parental NFL clones (Fig. 2A and C, compare lanes 2 and 7, 3 and 8, 6 and 9).

### Endo H glycosidase reveals high mannose and complex glycan species

Above, we saw that I559P increased gp160m mobility. Possible explanations are i) impaired glycan processing and/or ii) “sequon skipping”, reducing the total number of glycans. Without the NFL linker, I559P mutants were also uncleaved, as was DS (Fig. 2, lanes 17 and 19). To further investigate, denatured Env from a subset of mutants was digested with endoglycosidase H (endo H) to selectively remove high mannose glycans, followed by SDS-PAGE-Western blot. Duplicate blots were probed with anti-gp120+gp41 MAb cocktail or anti-gp41 MAb cocktails (Fig. 3). SOS “parent” VLP gp120 was partially endo H-sensitive; gp41 was endo H-resistant (Fig. 3, lanes 1, 2, 9 and 10, red and green dots). Gp120 and gp41 bands were faint or absent in I559P and DS (Fig. 3, lanes 3-8 and 11-16). Instead, gp160m was dominant in the I559P mutant and, upon endo H treatment, migrated far more rapidly than parent gp160m (Fig. 3, lanes 2, 4, 10 and 12, magenta dots). In contrast, endo H-treated DS gp160m was a smear, like parent gp160m (Fig. 3, lanes 2, 6, 10 and 14). Endo H-treated double mutant gp160m showed an intermediate pattern (Fig. 3, compare even numbered lanes). All samples showed discrete, fully endo H-sensitive high mannose gp160i (Fig. 3, yellow dots). Overall, this confirms that I559P and DS mutants cause folding defects that result in poor gp160 processing. Furthermore, I559P increases gp160m’s endo H sensitivity, suggesting a prevalence of high mannose glycans.

**Figure 3.**
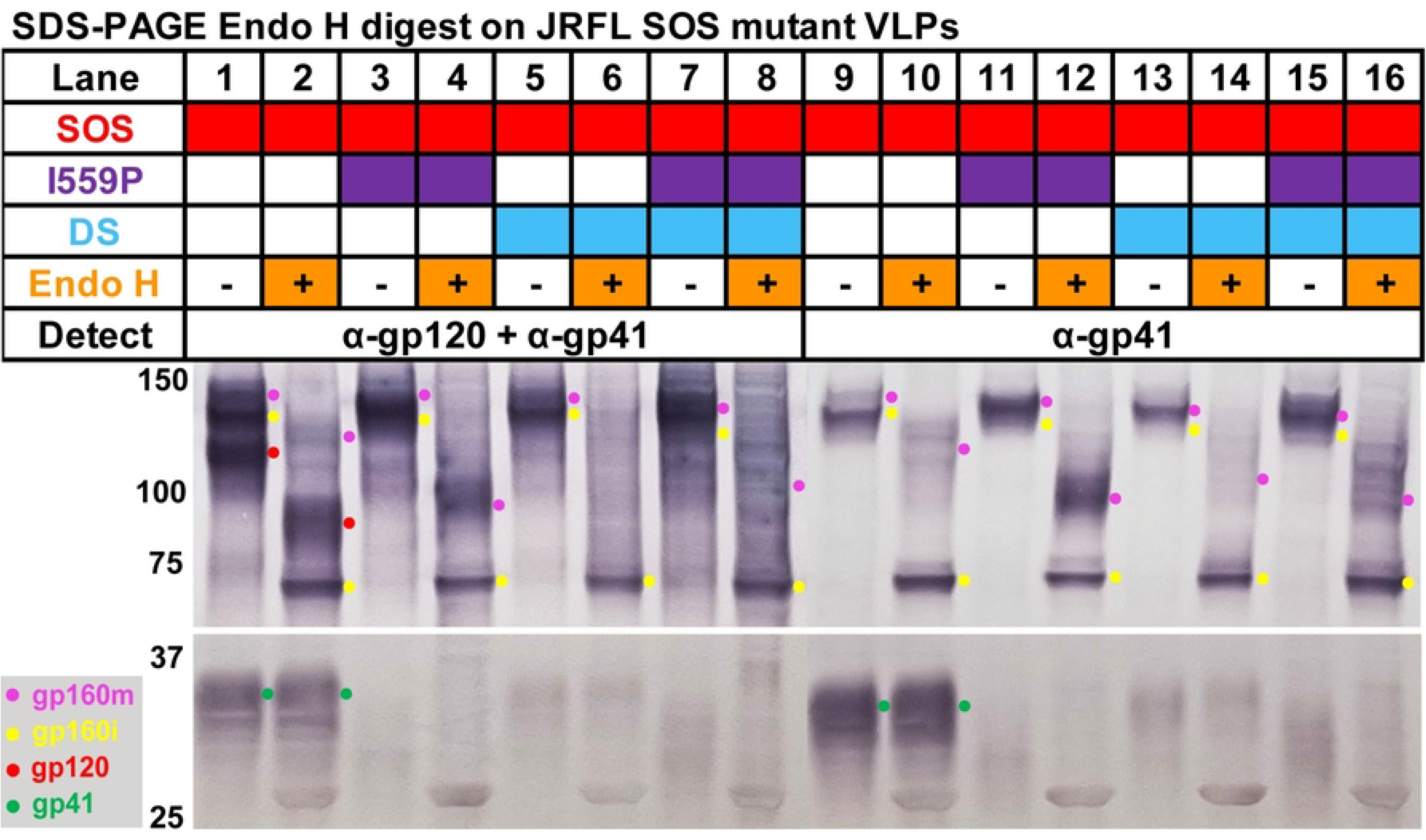
Endoglycosidase H digestion of I559P, DS and I559P+DS gp160ΔCT mutants. VLPs were lysed and boiled in SDS/DTT, then treated with endo H or PBS. Samples were then analyzed in duplicate SDS-PAGE-Western blots probed with anti-gp120+gp41 MAb cocktail (Lanes 1-8) or anti-gp41 MAb cocktail (Lanes 9-16).

### Enterokinase (EK) digestion generates gp120/gp41

An NFL linker-embedded enterokinase cleavage site (EK+) may provide a way to cleave gp120/gp41 post-expression [62]. We compared the EK sensitivity of NFL mutants with or without I559P and EK sites, i.e., samples from Fig. 2, lanes 2, 3, 7 and 8. EK digestion of EK+ but not EK-clones led to the appearance of gp120 and gp41 bands (Fig. 4A and B, compare lanes 11 and 15 to lanes 3 and 7; red and green dots, respectively). Concomitantly, gp160m was depleted (Fig. 4A and B, compare magenta dots in lanes 9, 11, 13 and 15). I559P did not affect gp160m depletion by EK digestion, (Fig. 4B, compare magenta dots in lanes 10 and 12, 14 and 16).

**Figure 4.**
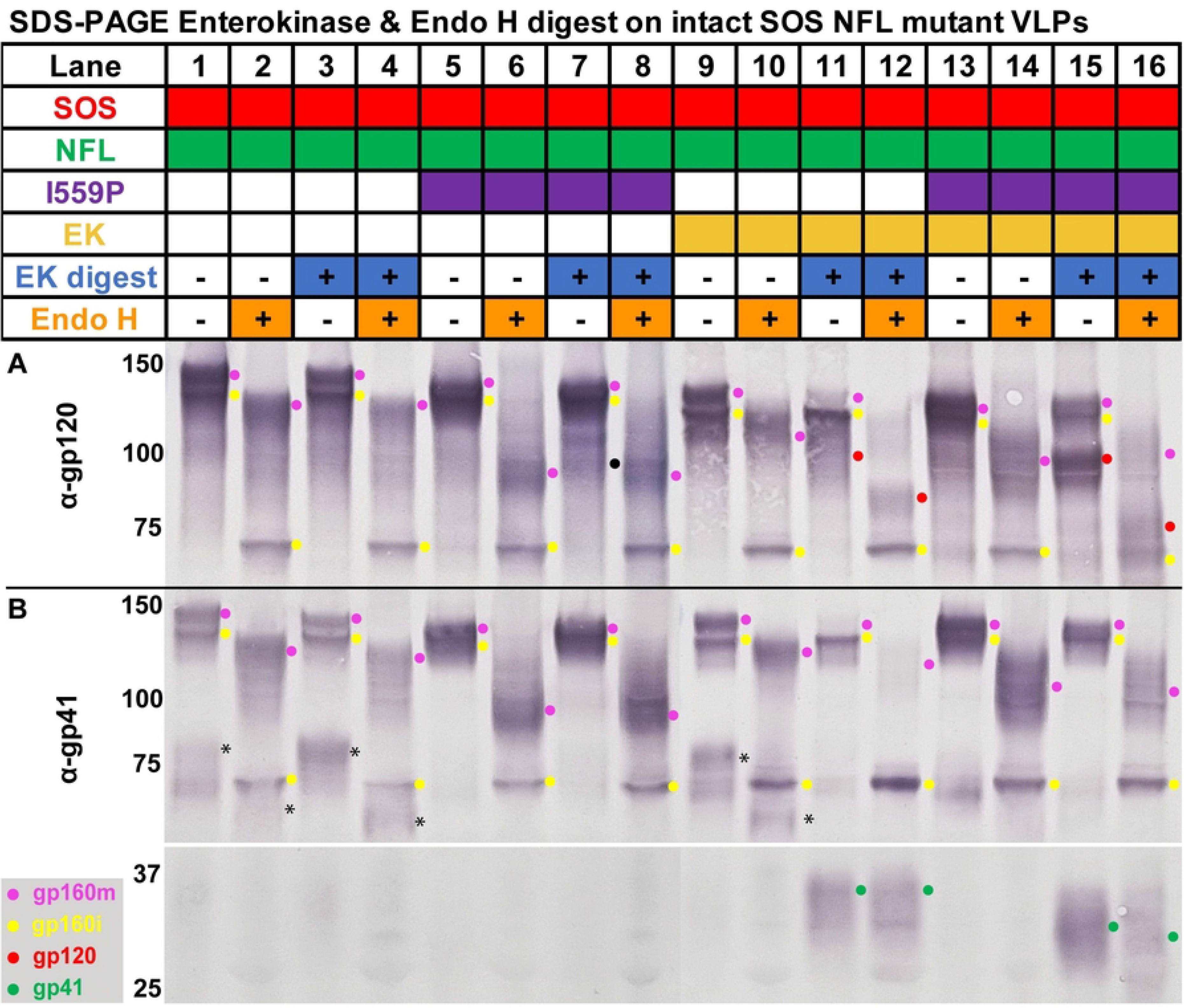
Enterokinase processing and endoglycosidase H digestion of gp160ΔCT SOS NFL mutants into gp120/gp41. VLPs were digested with enterokinase or PBS for 30h at 37°C, then lysed, incubated with endo H or PBS then analyzed in reducing SDS-PAGE-Western blot and probed with A) anti-gp120 or B) anti-gp41 MAb cocktails. The asterisk (*) marks an ∼80kDa band detected by anti-gp41 MAb cocktail in parent (non-I559P) samples, that suggests non-specific digestion, perhaps of the V3 loop, even in the absence of EK enzyme. The black dot detected in the anti-gp120 MAb cocktail (Lane 7) represents the non-specific digestion of gp160m by the EK enzyme.

As in Fig. 3, I559P gp160m bands migrated faster with endo H (Fig. 4A and B, compare even numbered lanes). EK-digested I559P gp120 and gp41 bands both migrated faster than their parental equivalents (Fig. 4A and B, compare lane 16 to lane 12). Indeed, I559P gp41 was partially endo H-sensitive, unlike the parent, suggesting modified glycan maturation (Fig. 4B, compare gp41 green dots in lanes 15 and 16 to lanes 11 and 12). We infer that I559P reduces glycan maturation in *both* gp120 and gp41.

Despite lacking an EK site, I559P EK-gp160m was partially digested non-specifically such that a weak band corresponding to gp120 was detected (Fig. 4A, lane 7, black dot). Although gp41 was absent, an ∼80kDa band was detected by anti-gp41 MAb cocktail in parent (non-I559P) clones (Fig. 4B, black asterisks in lanes 1, 3 and 9). This also suggests non-specific digestion, perhaps of the V3 loop, even in mock incubations (i.e., no EK added). These bands were diffuse and only partially endo H-sensitive, suggesting that they derive from gp160m (Fig. 4B, black asterisks in lanes 2, 4 and 10). In stark contrast, gp160i was insensitive to EK digestion (Fig. 4 and B, yellow dots), suggesting that the EK site is sequestered. Overall, EK cleaves gp160m, but does not cleave gp160i.

### BN-PAGE reveals various forms VLP Env and their lateral stability

As we reported previously, SOS gp160ΔCT resolves as trimers and monomers in BN-PAGE [22,30] (Fig. 5A, lane 1). The trimer band includes functional, cleaved gp120/gp41, while the monomer comprises of gp160m and gp160i. SOS NFL also exhibited trimers and monomers. However, the trimer density was weaker (∼35%), and the monomer was stronger (∼160%) relative to SOS (Fig. 5A, compare lane 1 and 2; S1A and C Fig). This suggests that gp160 either fails to form trimers effectively, or that they are unstable upon detergent lysis and electrophoresis. In other words, gp120/gp41 cleavage promotes lateral trimer stability. Overlaying the I559P mutation increased both trimer and monomer staining (Fig. 5A, compare lanes 1-3; S1 Fig). A putative dimer was also present in all I559P and DS mutants (Fig. 5A, lanes 3, 6, 8, 9, 11, 12, 17, 18, and 19; S1B Fig). UFO mutants expressed poorly, consistent with SDS-PAGE above (Figs 2 and 5A, lanes 4 and 10). NFL A433P Env resembled the NFL parent, although the weak trimer band was more diffuse (Fig. 5A, compare lanes 2 and 5). When A433P was combined with I559P, a weak and fast-moving trimer was restored (Fig. 5A, compare lanes 3 and 6). EK+ versions of these mutants exhibited band patterns like their EK-equivalents, but expression was weaker (Fig. 5A, compare lanes 2 and 7, 3 and 8, 6 and 9).

**Figure 5.**
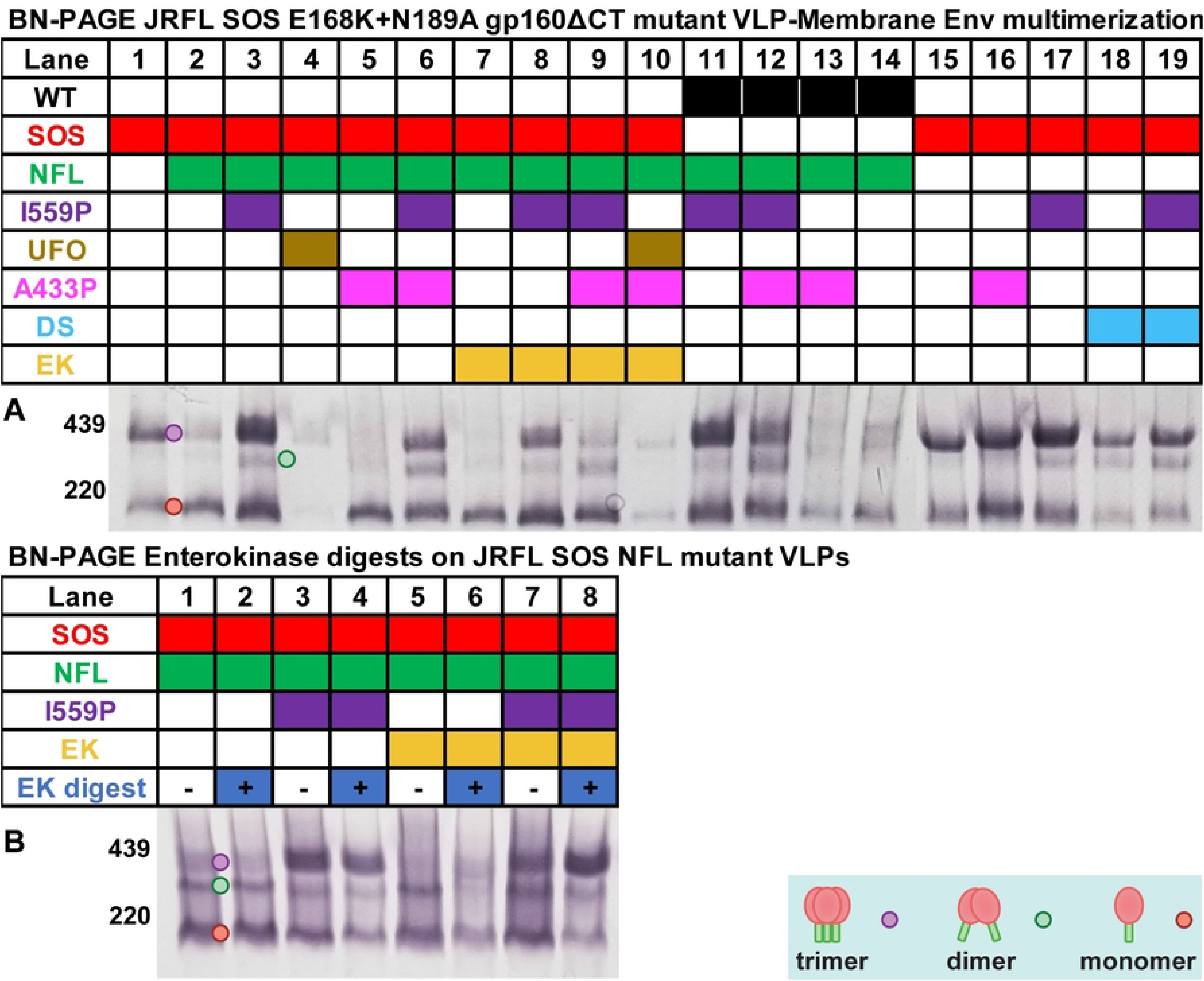
BN-PAGE-Western blot of gp160ΔCT mutants. A) The same mutants in Figure 2 were analyzed by BN-PAGE-Western blot, detecting with anti-gp120+gp41 MAb cocktail. Densitometry analysis of trimer, dimer and monomer band is shown in S1 Fig. B) BN-PAGE analysis of enterokinase-treated samples from Fig. 4. Env species are indicated by colored dots.

WT NFL I559P expressed well, with dimers like its SOS equivalent (Fig. 5A, lanes 3 and 11; S1 Fig). Overlaying A433P reduced Env expression, like its SOS counterpart (Fig. 5A, compare lanes 3 and 11 to lanes 6 and 12). WT NFL A433P expressed poorly and was diffuse, again like its SOS analog (Fig. 5A, lanes 5 and 13). WT NFL also expressed weakly, with diffuse bands (Fig. 5, lanes 2 and 14).

We next evaluated SOS clones without NFL. SOS A433P exhibited trimers and monomers, like the SOS parent (Fig. 5A, lanes 15-16), although monomer and trimer mobility were reduced. I559P exhibited predominant trimer, modest monomer and a faint dimer (Fig. 5, lane 17). DS exhibited weak trimer and monomer and a faint dimer (Fig. 5, lane 18). Combining DS and I559P resulted in intermediate staining (Fig. 5A, compare lanes 17-19). Overall, we infer that I559P improves expression and that lateral trimer stability is attained either by gp120/gp41 cleavage or by I559P mutants.

### Enterokinase (EK)-mediated gp120/gp41 processing increases lateral trimer stability

Since gp120/gp41 cleavage increases lateral stability, we investigated if post-expression EK-mediated cleavage of EK+ NFL gp160ΔCT into gp120/gp41 affects the trimer in BN-PAGE. We treated clones in Fig. 5A lanes 2, 3, 7 and 8 with or without EK enzyme (Fig. 5B). Mock treatment led to the appearance of dimers in non-I559P samples (Fig. 5B, lanes 1 and 5) that were absent in the corresponding untreated samples (Fig. 5A, lanes 2 and 7). EK digestion of EK+ clones led to changes in band patterns; I559P showed more trimer and less monomer (Fig. 5B, lanes 7-8). EK digestion of the non-I559P EK+ clone led to weaker monomer and dimer (Fig. 5B, lanes 5 and 6). In contrast, digestion of the EK-samples had little or no effect (Fig. 5B, lanes 1-4).

### NT1-5, UNC, A328G+/-D197N effects on membrane trimer

The NT1-5 combination mutant (M535I+L543Q+N553S+Q567K+G588R), which affects the “b” positions of the highly conserved gp41 N helix, was reported to reduce gp160 monomer expression, without affecting trimer expression [17,20]. Here, we evaluated the effects of NT1-5 in the context of JR-FL SOS gp160ΔCT. To enable us to better judge the effects of NFL, we also made a double mutant (K510S+R511S) [65], to create a simple uncleaved (UNC) mutant.

In SDS-PAGE, NT1-5 Env showed slightly reduced gp120/gp41 processing (S2A Fig, lanes 1, 2, 4 and 5). As expected, UNC lacked gp120 and gp41, and showed a strong gp160m band (S2A Fig, lanes 3 and 6). In BN-PAGE, NT1-5 increased trimer and decreased monomer (S2B Fig, compare lanes 1 and 2). In contrast, UNC was predominantly monomer, with weak dimer and trimer (S2B Fig, lane 3). This pattern resembles NFL (Fig. 5A, lane 2), except that the dimer is more prominent for UNC. Overall, this data suggests that, like I559P, NT1-5 laterally stabilizes UNC trimers and modestly reduces gp120-gp41 processing.

Previously, we used the globally sensitive A328G mutant to measure non-neutralizing antibodies in vaccine sera [26]. A328G was sensitive to non-NAbs 39F and 15e (S2C Fig, left panels, open symbols). Interestingly, D197N was somewhat b12-resistant compared to the parent but became more b12-sensitive when combined with A328G mutation. This overt change was mirrored by higher sensitivity to 39F and 15e compared to the other A328G mutant. In addition, PG16 sensitivity was lost, consistent with a “globally sensitive” conformation. In BN-PAGE, A328G showed reduced trimer, suggesting lower stability (S2D Fig, bottom panel, lanes 2 and 4). In SDS-PAGE, all clones showed very similar patterns (S2D Fig, top panel). We therefore chose SOS E168K+N189A+D197N+A328G as a prototype globally sensitive mutant for further experiments.

### BS^3^ crosslinking effect on membrane trimer

Although SOS trimers are stabilized in the apical plane by a disulfide bond, they may dissociate laterally following lysis and BN-PAGE [30]. To check this point, we used BS^3^ to crosslink selected clones. Crosslinking significantly reduced monomer in all cases, particularly for I559P or DS (Fig. 6, compare lanes 6, 8, 10, 12 to lanes 5, 7, 9, 11). Notably, SOS NFL I559P behaved similarly to SOS I559P (Fig. 6, lanes 5, 6, 9 and 10). Since I559P prevents gp120/gp41 cleavage, these clones differ only by the length of the peptide linking gp120 and gp41, which evidently has little effect here. For non-I559P mutants, some monomer shifted upwards slightly with BS^3^ (Fig. 6, lanes 2 and 4, black asterisks). The survival of some trimer in the face of boiling in SDS and DTT verifies that BS^3^ crosslinking was effective (Fig. 6, lanes 13 and 14). Overall, this is consistent with the idea that Env is mostly if not all trimeric but different forms can exhibit differing lateral stabilities.

**Figure 6.**
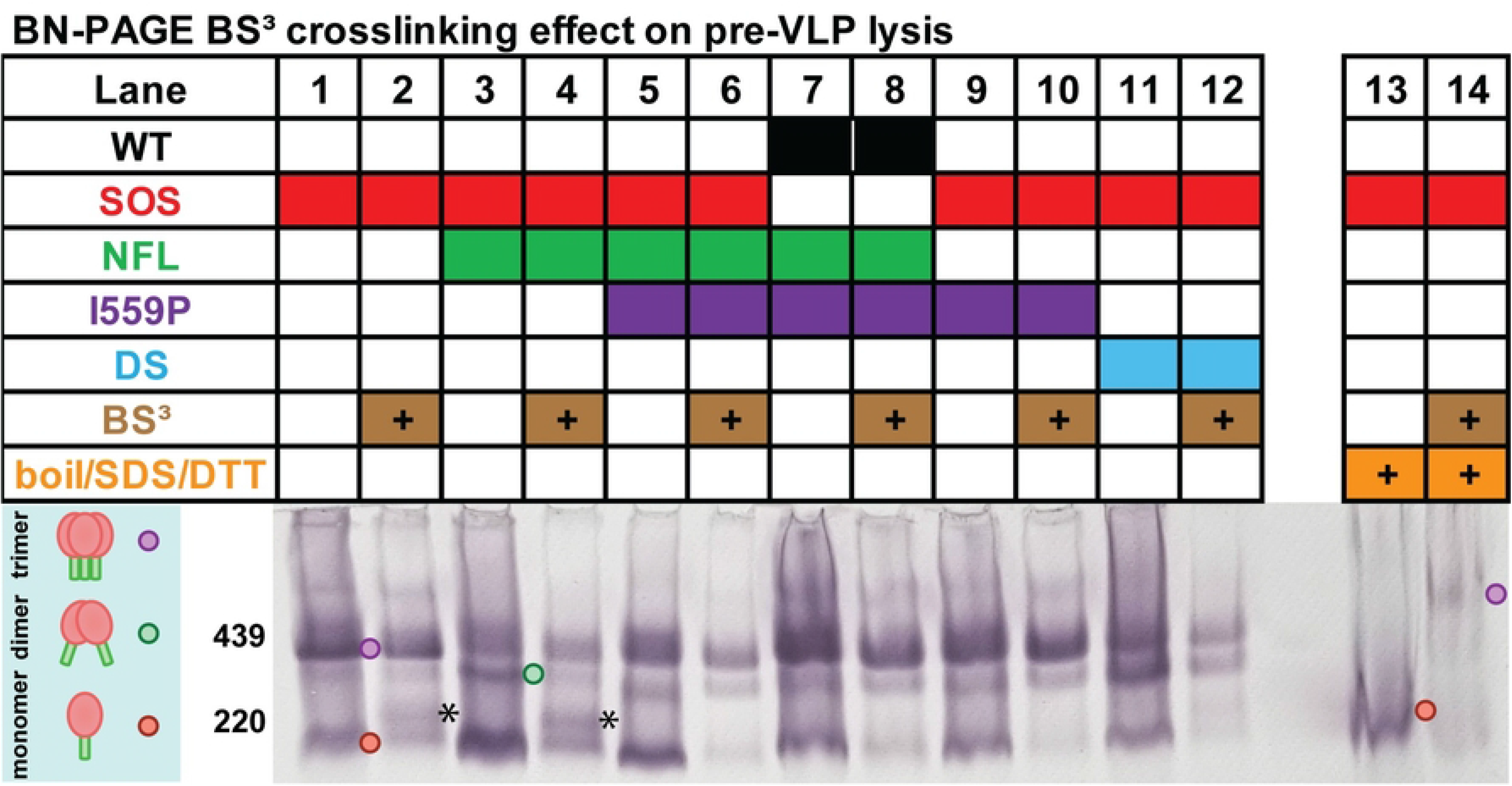
Effect of lysis on gp160ΔCT trimer stability in the presence of BS^3^ crosslinker. VLPs corresponding to lanes 1, 2, 3, 11, 17 and 18 of Figs 2 and 5 were crosslinked with and without BS^3^, then analyzed by BN-PAGE-Western blot, detecting with anti-gp120+gp41 MAb cocktail. Crosslinking efficiency was checked by boiling crosslinked or PBS-treated SOS VLPs in SDS and DTT (lanes 13 and 14). The asterisk (*) marks monomer shifted upwards with BS^3^ crosslinking. Env species are indicated by colored dots.

### Effect of DS on membrane trimer-sCD4 sensitivity

The DS mutant was designed based on the SOSIP gp140 trimer structure to insert a disulfide between positions 201 and 433 to restrict soluble CD4 (sCD4) binding and downstream conformational changes [11]. Soluble DS gp140 trimers bind only one molecule of sCD4, preventing full exposure of CD4-inducible (CD4i) and V3 epitopes. To check this with membrane trimers, excess 4 domain sCD4 was incubated with parent or DS VLPs, followed by a wash and BN-PAGE to evaluate shifts. DS trimers were shifted like the parent trimers, suggesting the binding of 3 sCD4 molecules (S3 Fig, compare lanes 2 and 4) [30]. Overall, the similar sCD4 binding patterns suggest that DS does not limit sCD4 binding to membrane trimers.

### Probing VLP antigenicity by native PAGE

Authentic, functional trimers are likely to be important for vaccine efficacy. A key question is whether trimer stabilizing mutations have unwanted antigenic consequences. Flow cytometry and cell ELISA have been used to appraise binding of NAbs and non-NAbs [10,29]. However, these methods can not differentiate between binding to native trimers or other forms of Env presented on cell surfaces, including gp160m. “BN-PAGE shifts” [30] can provide some clarity on this point by revealing MAb complexes with different Env isoforms. We first compared MAb binding to SOS and SOS NFL I559P trimers (Fig. 7) using a panel of NAbs directed to CD4bs, V2 and interface epitopes. Non-NAbs directed to the V3 and CD4bs were included to monitor unwanted epitope exposure. To detect MAb-Env complexes, blots were initially probed directly with anti-human IgG AP conjugate (Fig. 7B and D). Conversely, the blots in Fig. 7A and C were probed with anti-Env cocktails to detect *all* forms of Env, regardless of whether it is uncomplexed or complexed with MAb.

**Figure 7.**
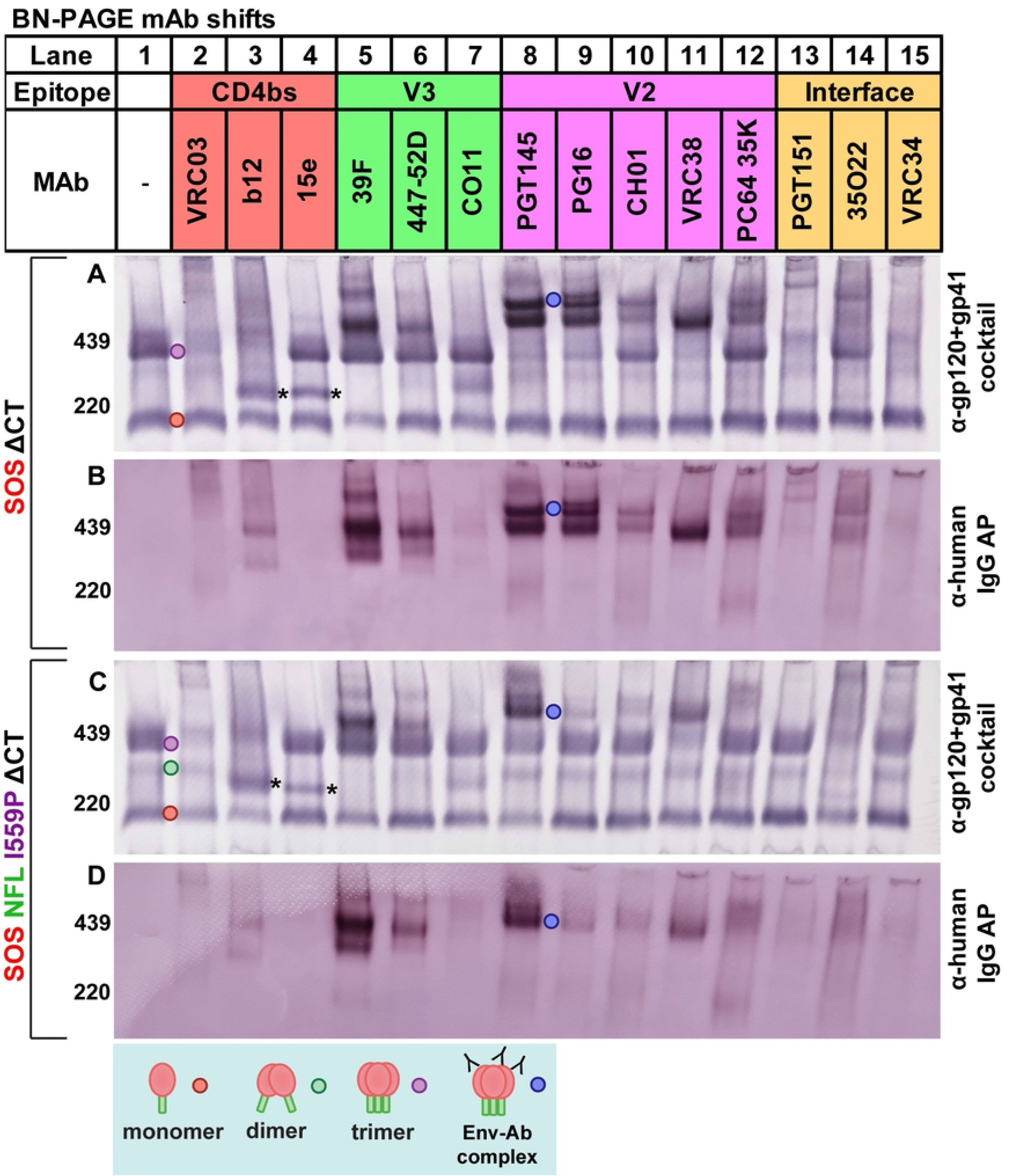
MAb binding to JR-FL gp160ΔCT E168K+N189A in BN-PAGE “shifts”. MAbs were mixed with SOS VLP (A and B) or SOS NFL I559P VLP (C and D) and were incubated at 37°C for 1h, then washed, lysed and analyzed by BN-PAGE-Western blot, probing with anti-gp120+gp41 MAb cocktail, followed by anti-human IgG AP conjugate (A and C) or just the anti-human IgG AP conjugate (B and D). The asterisk (*) marks a ∼220kDa observed with b12 and 15e, possibility arise from decreased lateral stability during MAb binding.

### CD4bs MAbs

VRC03 and b12 depleted uncomplexed SOS or SOS NFL I559P trimers (Fig. 7A and C, compare lanes 1, 2 and 3). In contrast, 15e did not bind to either trimer (Fig. 7A and B, lane 4). ∼220kDa bands were observed with b12 and 15e (Fig. 7A and C, lanes 3 and 4 asterisks). There were no bands in lanes 3 and 4 of Fig. 7B and D that could account for the ∼220kDa bands in the corresponding lanes Fig. 7A and C (see asterisks in Lanes 3 and 4), suggesting that these are ‘Env only’ bands, not MAb-Env complexes. Notably, the monomer was only partially depleted at best by MAb binding.

### V3 MAbs

39F, 447-52D and CO11 did not significantly deplete either trimer (Fig. 7A and C, compare lane 1 to lanes 5-7). 39F and 447-52D induced bands at ∼440kDa. The same bands were present in the corresponding “conjugate alone” blots (Fig. 7A-D, lanes 5-6), suggesting that they are MAb-Env complexes. In contrast, CO11 induced a ∼300kDa band (Fig. 7A and C, lane 7). To try to understand the ∼440kDa complexes induced by 39F and 447-52D, we evaluated 39F shifts of the SOS NFL (Fig. 8A, lane 2). Although SOS NFL resolves mostly as a monomer in BN-PAGE, 39F still induced a ∼440kDa species, indicating that 39F binds to uncleaved trimers and prevents them from dissociating laterally (Fig. 8A, lane 1).

**Figure 8.**
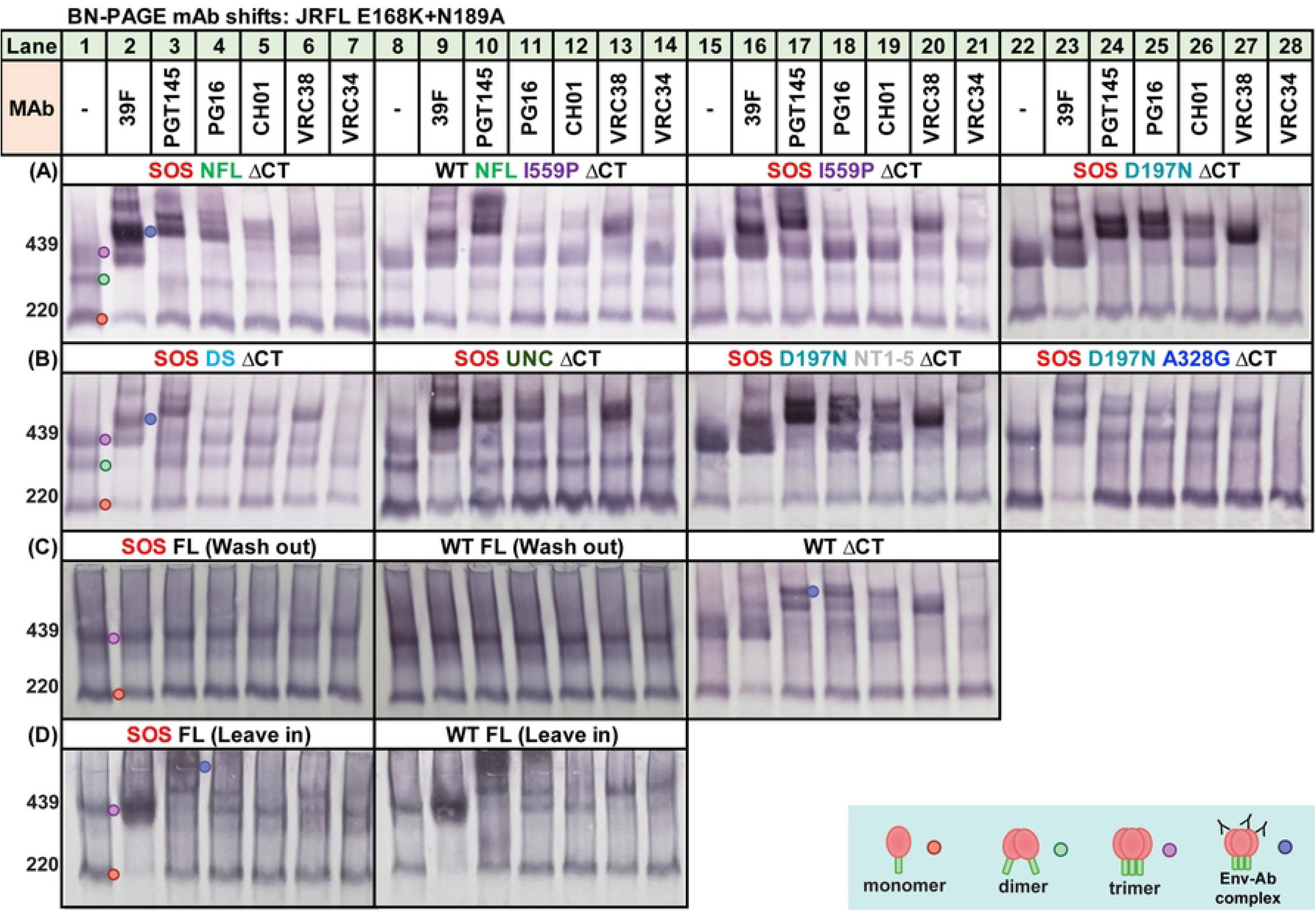
Dissecting mutant contributions to antigenic profiles by BN-PAGE mAb “shifts”. MAb shifts were performed with various mutants and resolved on BN-PAGE, detecting with anti-gp120+gp41 MAb cocktail. Mutants are all JR-FL gp160ΔCT E168K+N189A, except for some full length (FL) gp160 clones. (A) SOS NFL (lanes 1-7), WT NFL I559P (lanes 8-14), SOS I559P (lanes 15-21), SOS D197N (lanes 22-28); (B) SOS DS (lanes 1-7), SOS UNC (Lanes 8-14), SOS D197N NT1-5 (lanes 15-21), SOS D197N+A328G (lanes 22-28); (C) SOS FL – Wash out (lanes 1-7), WT FL – Wash out (lanes 8-14), WT (lanes 15-21); (D) SOS FL – Leave in (lanes 1-7), WT FL – Leave in (lane 8-14). For VLPs bearing FL Env, excess MAbs are either removed by washing in PBS (Wash out) or not removed (Leave in). UNC mutant is gp160ΔCT SOS E168K+K510S+R511S.

### V2 MAbs

PGT145, PG16, CH01, VRC38 and PC64 35K all bound to both SOS and SOS NFL I559P (Fig. 7A and C, lanes 8-12), albeit with differences. SOS trimers were completely depleted by PGT145 and PG16 (Fig. 7A and C, lanes 8-9). PGT145, PG16, CH01 and PC64 35K shifts of SOS Env resulted in two high molecular weight bands at ∼440kDa and ∼480kDa (Fig. 7A, lanes 8, 9, 10 and 12). However, for SOS NFL I559P, only a ∼440kDa band was clear with PGT145 (Fig. 7C, lane 8). VRC38 also shifted SOS more completely than SOS NFL I559P (Fig. 7A and C, lane 11). Indeed, V2 NAb binding to SOS NFL I559P was generally weak: trimer depletion was marginal and shifted bands were fainter (Fig. 7C, lanes 8-12). The same blots probed with only anti-IgG AP conjugate (Fig. 7B and D, lanes 8-12) add further support weaker V2 NAb binding to SOS NFL I559P, contrasting sharply with the equivalent binding of 39F and 447-52D to uncleaved Envs of the two mutants (Fig. 7B and D, lanes 5 and 6).

To better understand PGT145 binding patterns, we examined intermediate mutants. The 440kDa and 480kDa doublet observed in Fig. 7A, lane 8 was prominent only in clones that permit at least some gp120/gp41 processing such as SOS D197N (Fig. 8A, lane 24), SOS D197N NT1-5 (Fig. 8B, lane 17) and WT ΔCT (Fig. 8C, lane 17). However, in those with poor or no processing (e.g., SOS NFL, WT NFL I559P, SOS I559P, SOS DS, SOS UNC), the 480kDa band was weak or absent. This suggests that PGT145 binding creates distinct complexes with cleaved gp120/gp41 trimers (480kDa band) and UNC trimers (440kDa band).

### Interface NAbs

PGT151 and VRC34 depleted SOS trimers but not SOS NFL I559P trimers, consistent with the importance of gp120/gp41 processing for their binding (Fig. 7A and C, lanes 13 and 15). In contrast, 35O22 depleted SOS NFL I559P trimers more effectively than SOS trimers (Fig. 7A and C, lane 14). Since 35O22 binding is subject to glycan clashes [46], the improved binding may reflect glycan changes associated with I559P. We revisit this point below.

### Other NAbs

Lastly, we further evaluated the two main mutants of Fig. 7 in BN-PAGE shifts with a broader panel of MAbs (S7 Fig). As expected, 2G12, PGT121, 2F5 and 4E10 exhibited comparable binding patterns with both mutants. Neither 7B2 nor CR3022 bound. In the case of 7B2, the SOS mutation occludes the gp41 cluster I epitope, while lack of CR3022 binding confirms specificity.

### Intermediate mutants reveal the dominant effect of I559P on Env antigenic profile

We further investigated antigenic differences using a focused MAb panel with intermediate mutants. SOS NFL was more V2-sensitive than SOS NFL I559P (compare Fig. 7C to Fig. 8A, lanes 3-6), further emphasizing the adverse effect of I559P on V2 epitopes. Accordingly, WT NFL I559P (Fig. 8A, lanes 10-13) and SOS I559P (Fig. 8A, lanes 17-20) showed only modest V2 NAb binding. Overall, V2 NAb binding to I559P-bearing mutants was weak, particularly for PG16 and CH01. Neither SOS nor NFL contributed to this effect. Binding of V2 MAbs to SOS DS and D197N NT1-5 was also weak: like I559P mutants, PG16 and CH01 were markedly affected (Fig. 8B, lanes 4, 5, 18 and 19). In contrast, like the SOS parent, SOS D197N (Fig. 8A, lanes 25 and 26) showed robust V2 NAb binding; PG16 and CH01 depleted the uncomplexed trimer effectively. VRC34 binding to I559P mutants was also weak (Fig. 8A, lanes 14 and 21). In contrast, 39F binding to all mutants was comparable, except for full-length (FL) clones in the wash out method (covered below). Notably, SOS I559P (Fig. 8A, lanes 15-21) is a close relative of SOS NFL I559P (Fig. 7C), both being uncleaved and differing only in terms of the linker (UNC vs NFL). This difference did not register in BN-PAGE shifts: V2 NAb binding was poor in both cases. WT ΔCT exhibited a very similar pattern to SOS ΔCT (compare Fig. 7A to Fig. 8C, lanes 15-21), suggesting that SOS exerts only mild antigenic effects. Globally sensitive SOS D197N+A328G exhibited reduced V2 NAb binding (Fig. 8B, lanes 24-27), consistent with the expected loss of these epitopes.

### Analysis of full-length gp160 VLPs

Much of our prior vaccine development work has used gp41 truncated Env (ΔCT), due its improved expression, gp120/gp41 processing and minor effects on antigenic profile [22]. However, in some Env strains, SOS and ΔCT mutations can affect antigenic profile [36,38,66,67]. We therefore evaluated WT and SOS JR-FL clones in ΔCT and gp160 full-length (FL) formats in SDS-PAGE with or without endo H, then probed in duplicate blots with gp120 and gp41 MAb cocktails. FL clones exhibited very weak gp120/gp41 processing, revealed by faint endo H-resistant gp41 bands (S4A Fig, lanes 9-16). Corresponding gp120 bands were also weak (S4A Fig, compare lanes 3, 4, 7 and 8 to 1, 2, 5 and 6). As expected, gp160 and gp41 bands in FL clones migrated slower due to the additional gp41 mass. Notably, FL clones exhibited very strong endo H-sensitive gp160i bands (S4A Fig, compare yellow dots in lanes 3 and 7 with 1 and 5), suggesting an elevated rate of misfolding.

In BN-PAGE shifts, MAb-VLP complexes were washed to remove unbound MAb prior to lysis and electrophoresis. Intriguingly, in this format, both FL clones *did not show binding by any MAb* (Fig. 8C, lanes 1-14). However, binding was observed *when the wash step was omitted*. 39F robustly shifted the WT and SOS FL monomer (Fig. 8D, lanes 2 and 9). FL shift patterns (Fig. 8D, lanes 1-14) resembled ΔCT counterparts (Fig. 7A and 8C lanes 15-21). However, VRC38 and VRC34 binding to WT FL trimer was stronger than to SOS FL trimer (Fig. 8D, compare lanes 6-7 and lanes 13-14) or SOS ΔCT mutant (Fig. 7A, lanes 11 and 15). Furthermore, CH01 showed little or no binding on FL clones and PG16 was only partially saturating.

Two possibilities may account for the lack of MAb binding to FL in the washout format. First, FL undergoes a conformational change when liberated from membranes that results in MAb dissociation, or second, none of these MAbs bind to FL trimers, so that binding observed in the “leave in” format occurs only *after* VLP lysis. To dissect these possibilities, we checked PGT145 in further BN-PAGE shifts in which BS^3^ was used to cross-link any PGT145-trimer complexes to prevent MAb dissociating upon lysis (S5 Fig). In addition to comparing the effects of washing after PGT145 incubations, we also compared the effects of washes after BS^3^ crosslinking. We found that PGT145 failed to shift trimer in any format that included a wash, even when BS^3^ was added (S5 Fig, compare lanes 7 and 12). In the leave in format, there was an additional low molecular weight band when BS^3^ was added, without subsequent washing (S5 Fig, lane 8). This band was not observed when PGT145 was omitted (S5 Fig, lane 6), so this could be free MAb that pellets with VLP due to non-specific cross-linking but is not bound to Env. Taken together, this data supports that the second possibility, that MAbs simply do not bind to FL on intact VLPs and can only bind *after* lysis.

We next explored the effects of SOS and ΔCT mutations in another strain, PC64 MRCA from a subtype A-infected donor [68]. SDS-PAGE blots revealed smeary gp160 and gp120 bands in ΔCT format but tighter bands in FL format, suggesting more uniform glycans (S4B Fig, compare lanes 1-4). Like JR-FL ΔCT, PC64 ΔCT exhibited far more efficient gp120/gp41 processing than its FL counterpart. In BN-PAGE, SOS appeared to improve trimer and monomer expression relative to WT (regardless of ΔCT or FL), and ΔCT clones were better expressed (S4C Fig). Gp120 shedding was detected for WT ΔCT, resulting in gp41 stumps (S4C Fig, compare lanes 3 to lane 1).

The neutralization sensitivity profiles of PC64 WT ΔCT and FL clones were similar (S6 Fig). The SOS FL clone was insufficiently infectious and was therefore omitted from this analysis.14E sensitivity was modestly increased by SOS in ΔCT format, as was b12 sensitivity. Correspondingly, PG16 sensitivity was reduced. Collectively, this suggests that SOS converts the trimer into a globally sensitive phenotype. All clones were CH01-resistant. Together, this shows that FL clones exhibit weaker gp120/gp41 processing and expression. More importantly, the effects of SOS on NAb sensitivity suggest a greater conformation change than observed for our JR-FL clones. Therefore, some effects of mutations are universal (expression and cleavage), but others (NAb sensitivity) are context dependent.

### Probing VLP antigenicity by virus capture

To further probe the antigenic effects of modifications, we tested the ability of immobilized MAbs to capture VLPs. SOS VLPs were effectively captured by 2G12, PGT145, PG16 and VRC38. CH01 and PC64 35K were less effective among the V2 mAbs. 39F and 447-52D efficiently captured SOS VLPs, while CO11 capture was poor (S8A Fig). This is consistent with BN-PAGE shifts, where 39F and 447-52D bound more effectively (Fig. 7B). On the other hand, these V3 MAbs neither bound to native trimer (Fig. 7A), nor neutralized (S8B Fig). PGT121 capture was highly effective (S8A Fig). VRC01, VRC03, b12 and 15e all captured effectively (S8A Fig). 15e also captured well without neutralizing. PGT151, 35O22 and VRC34 captured moderately. 10e8, but not 2F5, captured well. 7B2 and 2.2B both captured poorly, as the SOS mutation eliminates these epitopes [30]. Overall, most NAbs captured with at least modest efficiency. Non-NAbs variably captured, presumably via binding to UNC.

Previously, we found that JR-FL WT and SOS PVs exhibited similar neutralization profiles, as did ΔCT [61]. However, this comparison did not include the various ‘new wave’ of bNAbs reported since 2009. We therefore compared NAb sensitivity profiles of WT, SOS, FL and ΔCT using a comprehensive set of MAbs (S8B Fig). SOS FL infectivity was weak, (S9 Fig), and was therefore excluded from NAb sensitivity profiling. The 3 other clones exhibited largely consistent profiles (S8B Fig). However, SOS gp160ΔCT was more sensitive to 2G12, b12, 10E8 and 2F5 compared to WT gp160ΔCT. The additional sensitivity to MPER MAbs afforded by the disulfide bond that must be broken by exposure to reducing agent, providing more time for these MAbs to neutralize.

Since the 2G12 epitope is constitutively exposed [10,37,40], relative 2G12 capture can be used to standardize capture for each Env clone, regardless of any expression differences. There was a ∼2-fold range of 2G12 capture efficiency, with SOS NFL A433P being most sensitively captured, while both WT and SOS FL were the least well captured (S10A Fig).

We next evaluated JR-FL mutant MAb capture, expressed as % relative to 2G12 capture (Fig. 9). Capture of each VLP clone was compared to SOS-VLPs, testing for significant differences by Kruskal-Wallis non-parametric rank test. MAb capture of WT ΔCT and SOS ΔCT was similar. Modest differences in PGT145, PG16 and CH01 capture were not significant. Conversely, VRC38 capture of WT FL was significantly weaker. 15e and 39F capture of FL clones was weaker than their ΔCT counterparts. Next, we consider MAb capture of four SOS ΔCT variants: SOS (parent), SOS NFL I559P, SOS D197N+NT1-5 and SOS D197N. PG16 and CH01 captured SOS NFL I559P less effectively, consistent with weaker BN-PAGE shifts (Fig. 7).

**Figure 9.**
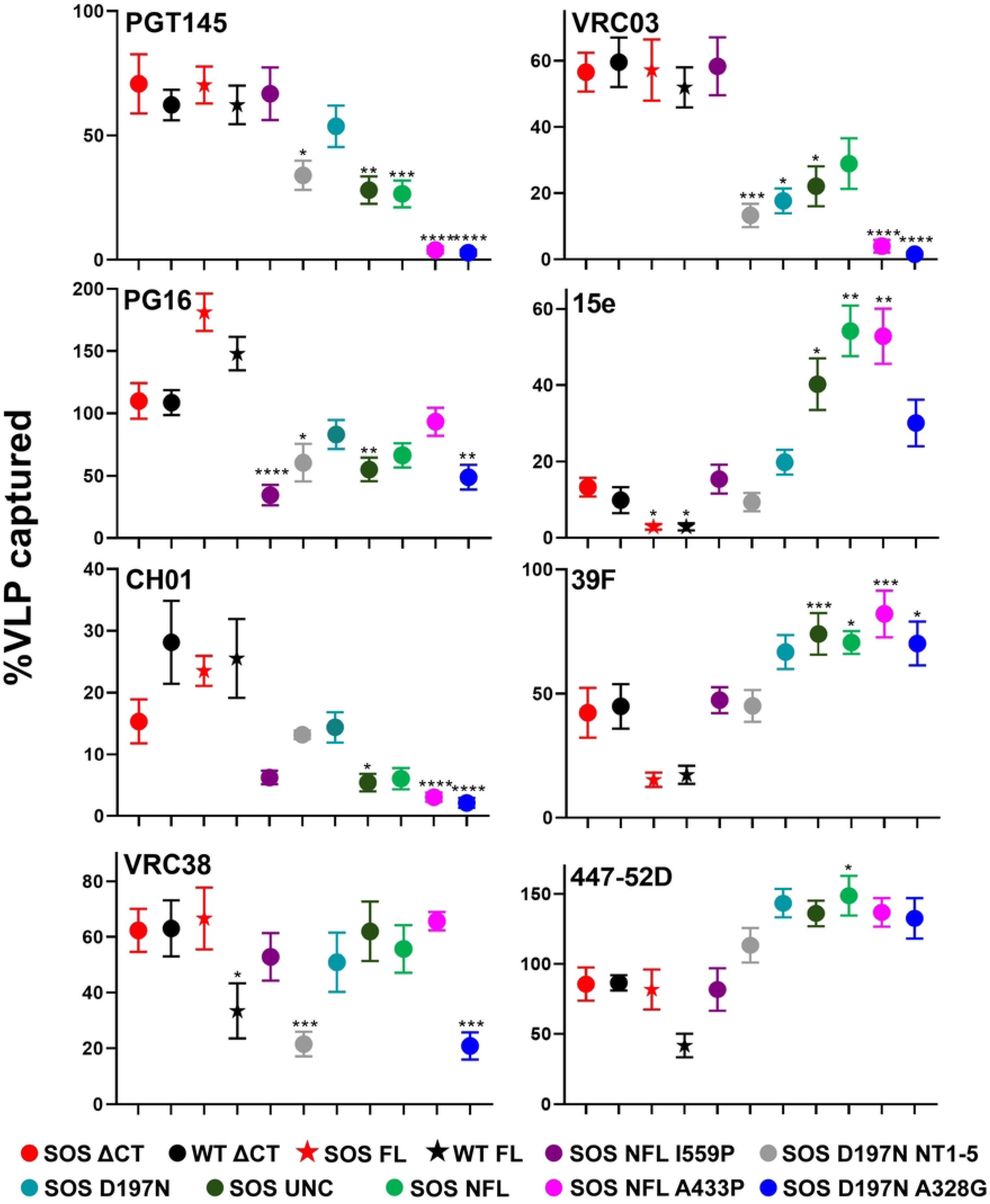
Antigenic profile of VLPs carrying different mutations assessed by virus capture assay. Capture of various JR-FL mutant VLPs by a subset of MAbs. For each mutant, data was normalized by 2G12 capture which approximates Env expression. This allows mutants to be compared regardless of expression differences. Virus capture assays were performed in quadruplicate and repeated at least three times. Error bars represent the standard deviation of the mean. UNC is gp160ΔCT SOS E168K+K510S+R511S. Kruskal-Wallis test was used to analyze for significant difference of different VLPs captured by a MAb. \**p <* 0.05, ** *p* < 0.01, *** *p* < 0.001, **** *p* < 0.0001.

Previously, NT1-5 reduced non-NAb capture, but did not affect NAb capture [17]. However, here we observed that D197N+NT1-5 was captured poorly by all V2 MAbs (Fig. 9), while D197N mutant alone did not significantly affect capture, although it trended to weaker capture. This reflects the poor V2 binding to NT1-5 by BN-PAGE (Fig. 8). Overall, we infer that the I559P and NT1-5 both disrupt V2 epitopes. VRC03 capture was equivalent for SOS and SOS NFL I559P but was close to background for the D197N-containing mutants. This suggests that the N197 glycan clashes with VRC03. Contrasting previous data [17],15e, 39F and 447-52D capture was comparable for all 4 mutants, although the D197N trended higher.

Next, we considered UNC and NFL. PGT145 only modestly captured both clones, suggesting that I559P is important for PGT145 binding to SOS NFL I559P (Fig. 9). PG16 and CH01 capture was also poor, like SOS NFL I559P. However, VRC38 capture was unaffected gp120/gp41 cleavage. VRC03 capture was weak, as its quaternary epitope is evidently poorly presented in uncleaved mutants. Conversely, 15e and V3 capture was effective. Overall, we conclude that I559P differentially affects V2 epitopes, with NFL having little impact.

PGT145, CH01 and VRC03 failed to capture SOS NFL A433P and SOS D197N+A328G mutants, consistent with their “globally sensitive” phenotypes. PG16 and VRC38 captured A433P moderately, but not D197N+A328G. Both mutants were well-captured by 15e, 39F and 447-52D, albeit to differing extents. Overall, we infer that both mutants adopt tier 1 conformations, but with distinct antigenic profiles [10,69]. Lastly, there was no detectable CR3022 capture, indicating the specificity of the assay (S10B Fig).

### I559P reduces HIV+ plasma recognition

In a previous study, two HIV-1 seropositive (HIV+) donors plasmas bound significantly more strongly to WT than SOS I559P membrane Envs [38]. We extended this analysis by testing the binding of 22 HIV+ plasmas from chronic infections [68,70] to JR-FL WT, SOS and SOS I559P VLPs by ELISA. Exemplary ELISA titers are shown in S11 Fig. 2G12 bound to all 3 Env VLPs equivalently, suggesting comparable expression. 2G12 did not bind to MuLV Gag “bald” VLPs (S11 Fig). A HIV-1-negative plasma, 210 showed only background binding, regardless of Env expression (S11 Fig). Plasma binding to SOS VLPs was weaker than to WT VLPs (Fig. 10 and S11 Fig). This may be due to the lack of immunodominant gp41 stumps on SOS VLPs. Weaker plasma binding to SOS I559P compared to SOS may be due to conformational differences, exemplified by reduced V2 bNAb reactivity to SOS I559P. Overall, the diminished recognition of SOS I559P versus SOS Env suggests I559P (Fig. 10) incurs conformational effects that reduces recognition by HIV-1+ plasma, including reduced recognition by NAbs.

**Figure 10.**
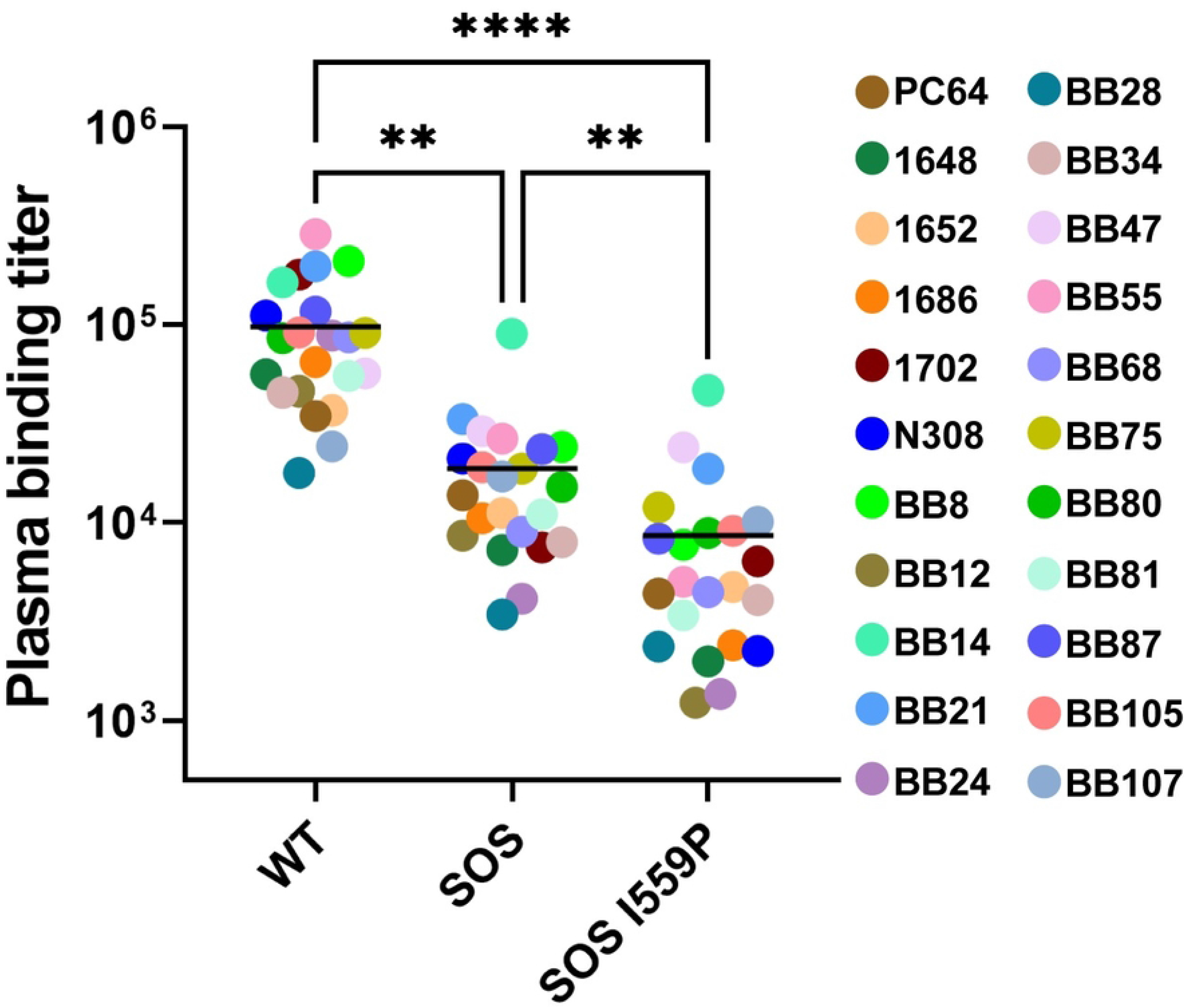
JR-FL gp160ΔCT E168K+N189A VLP binding titer of human HIV+ plasma probed by VLP ELISA. Plasma binding titer of human HIV+ plasma (n = 22) to JR-FL gp160ΔCT WT, SOS or SOS I559P VLP analyzed by ELISA. Plasma binding titers were calculated as the dilution where its binding OD was 0.5. Geometric mean binding titer (GMT) against each VLP is indicated as horizontal line. One-way ANOVA Dunn’s multiple comparison test revealed a significant difference of plasma binding titer between WT, SOS and SOS I559P VLPs. \*\**p* < 0.01, \*\*\*\**p* < 0.0001.

### Glycan analysis

To better understand the basis of the effects of JR-FL mutants, we assessed glycan occupation and maturation by glycopeptide in-line liquid chromatography mass spectrometry (LC-MS) [22,71,72]. In a previous study [22], we used this method to analyze unfractionated “total VLP” samples, in which the resulting data derives from the sum of various forms of Env that we described above in Western blots, including gp120/gp41 trimers, gp160m, gp160i and gp41 stumps. To try to generate “cleaner” samples for analysis and to enable us to discriminate between the glycan profiles of different Env isoforms, here, we ran VLP samples on reducing SDS-PAGE and cut out bands for analysis. In the case of the parent sample, we cut out 3 bands corresponding to gp160 (including gp160m and gp160i, which could not be separated sufficiently by this method), gp120 and gp41. For the other samples (NFL, NFL I559P and NFL A433P), we cut out gp160 bands (gp160m and gp160i), as there are no gp120 and gp41 bands when the NFL is used.

Detailed glycopeptide data is provided in S1 Raw glycopeptide folder, containing a file for each sample. Data was obtained for all samples except for the gp41 band of the parent VLP, which was insufficient for detection. Data is plotted in bar charts (S12 Fig) that reveal the relative percentages of oligomannose, hybrid, complex and core glycans at each site, as well as unoccupied sequons due to skipping. These charts also show fucosylated and sialylated glycans (NeuAc/NeuGc) at each site.

To summarize the data and to facilitate comparisons, each glycan type was given a score from 1 to 19, depending on the average maturation state (S1 Main glycan analysis). Specifically, the high mannose glycan, M9Glc, has a score of 1, while the most highly branched and fucosylated complex glycan HexNAc(6+)(F)(x) has a score of 19. Glycans at each site were then given an overall score based on the percentage of each glycoform at each site multiplied by its percent prevalence and rounded to the nearest whole number. Overall glycan scores were computed in S1 Main glycan analysis for each clone and are color coded for clarity. Glycan data were modeled for the gp160 bands (S13A Fig) and the gp120 band (S13B Fig) of the VLP parent. We also modeled “total VLP” parent samples (2020, 2021 and 2023) and gp120 monomer (2020) [22] (S13C, D, E and G Fig). Mutant glycan scores were also modeled (S14 Fig).

To determine the effects of mutations on glycan profiles, we first need to select a reference profile. The profiles of the 3 independent parent “total VLPs” (2020, 2021 and 2023) (S12, S13C-E, S1 Main glycan analysis score sheet) were largely similar. However, data for glycans at some positions were missing in some preparation but present in others. Therefore, we decided to combine and average the parent data as a reference to the mutants (S12 Fig, S13F Fig, S1 Main glycan analysis score sheet).

We next compared the glycan profiles of gp160 bands (containing gp160m and gp160i) and gp120 band with each other and to the average parent as well as the previous gp120 monomer (S13G Fig). Glycan score changes at each position are determined by subtraction. Score changes are color coded according to glycan processing: decreased processing is shown in shades of red and increased processing is shown in shades of blue (Fig. 11). For clarity, we refer to VLP gp120 as viral gp120 hereafter to distinguish it from gp120 monomer. There were stark differences between parent viral gp120 and gp160 bands, particularly at N356, that was completely flipped to complex in viral gp120. Increased viral gp120 glycan maturation was also found at N88, N188, N392 and N463. These are modeled in Fig. 11A, summarized in S1 Main glycan analysis and shown in detail in S1 Sheet.

**Figure 11.**
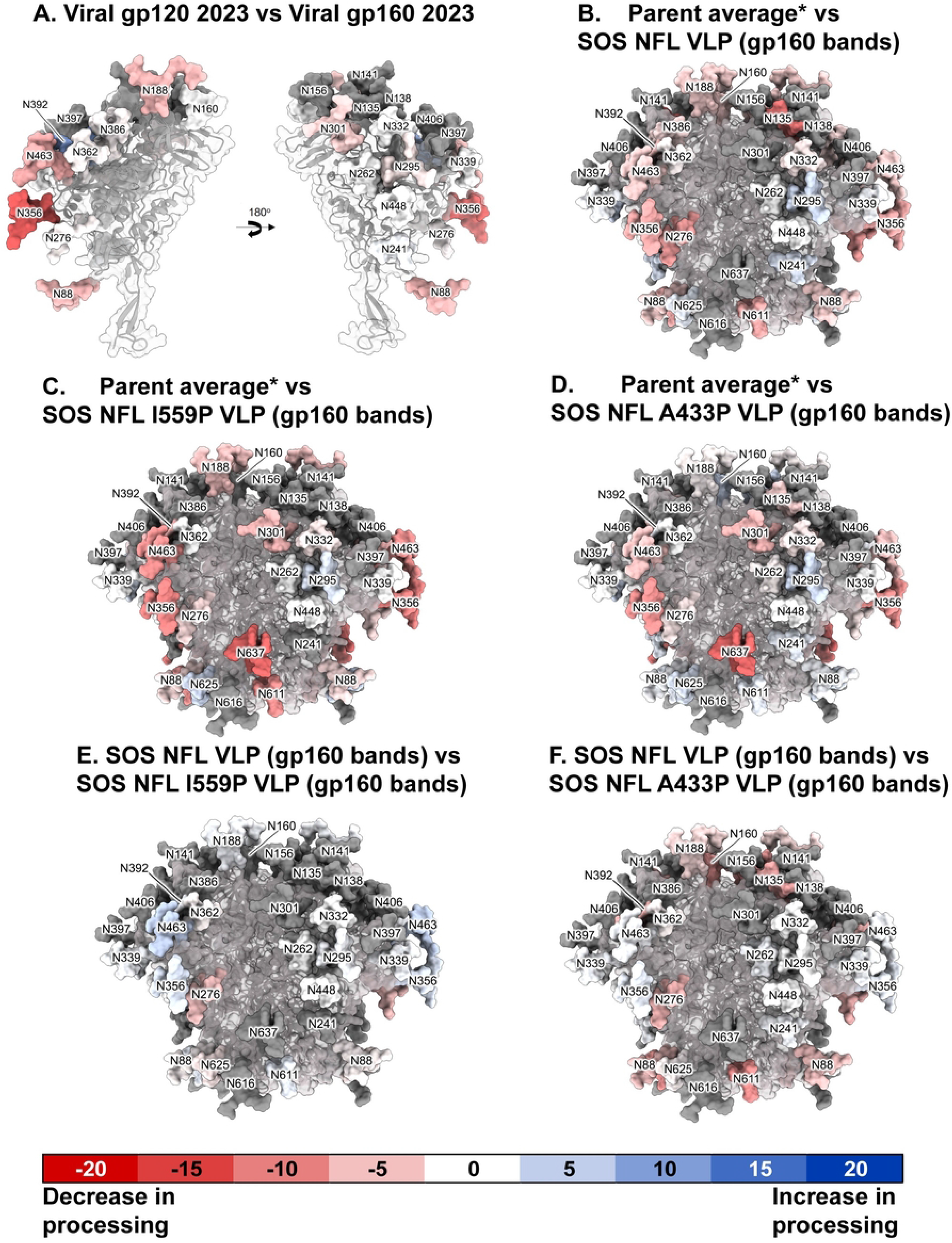
Effects of I559P and A433P mutants on gp160ΔCT glycan maturation and occupation. Related to S12-S14 Figs, S1 Main glycan analysis and S1 Raw glycopeptide folder. Each glycan on the trimer models (pdf: 6MYY) is numbered according to HxB2 strain and is given a maturation score, derived from LC-MS analysis. Glycan score differences between two clones are colored in shades of red (i.e. negative score indicates decrease in processing), white (no difference in glycan processing) or blue (i.e. positive score indicates increase in processing). Data are only shown at positions where a glycan was detected in >10% of the equivalent peptides of both samples in each pair. Some glycans, rendered in gray, were not resolved in the sample, and therefore have no score (not done; n.d.). Glycan score difference calculations between sample pairs are shown in S1 Main glycan analysis, and are modeled here as follow: (A) Viral gp120 band vs viral gp160 bands; (B) Parent average vs SOS NFL VLP gp160; (C) Parent average vs SOS NFL I559P VLP gp160; (D) Parent average vs SOS NFL A433P VLP gp160; (E) SOS NFL VLP gp160 vs SOS NFL I559P VLP gp160; (F) SOS NFL VLP gp160 vs SOS NFL A433P VLP gp160. * Parent average is the glycan score average of “total VLPs” preparation from 2020, 2021 and 2023 (S13F Fig).

Notably, viral gp120 did not show any sequon skipping. This contrasts with gp160 bands, where there was partial skipping at N188 and N339 (gray shading in S12A Fig, S1 Main glycan analysis). Total VLP samples also exhibited detectable skipping (S12A Fig). Mutant gp160s exhibited enhanced skipping, particularly at N637 (S12B Fig).

Comparing parent gp160 (S13A Fig) to parent average (S13F Fig) also revealed reduced glycan maturation at the same positions (S1 Sheet). In contrast, the glycan profiles of the parent average and the parent gp120 band (S13B Fig) were more consistent (S1 Main glycan analysis, S1 Sheet).

If viral gp120 and gp160m express equivalently, one might expect increased diversity at the variable glycan sites in total VLPs, as they would reflect the combined diversity of both gp120 and gp160m components (S1 Sheet). To some extent this is the case at N188 and N463 (especially the 2023 parent sample), but not so much at N88 and N356 that was biased more towards viral gp120 profile. In the 2023 total VLP sample, N392 exhibited intermediate processing. Overall, this suggests that functional gp120/gp41 trimers are the predominant form of Env in total VLPs and gp160m is the weaker form. Furthermore, the near complete complex glycans at some positions suggests that gp160m expression is dominant over gp160i, which exhibits high mannose glycans at all positions [46].

The gp120 monomer profile at variable positions N88, N160, N356 and N463 was also more akin to viral gp120 than the gp160 bands (S1 Sheet). However, the glycan at N392 was far more processed on gp120 monomer than on viral gp120 or total VLPs (S1 Sheet). We infer that N392 is uniquely buried in native trimers.

### Mutant glycan profiles compared to parent

The key information we sought here is whether mutants affect glycan compositions, as an indicator of conformational differences. We therefore compared the parent average to NFL, NFL I559P and NFL A433P (Fig. 11B-D, S1 Main glycan analysis). SDS-PAGE data above indicated that I559P bands were fast moving and slightly more endo H sensitive, whereas A433P bands were slightly slower moving. We hoped that glycan analysis would help us to reconcile these effects.

All 3 mutants trended towards reduced glycan processing (S1 Main glycan analysis score sheet). This implies that the simple lack of gp120/gp41 cleavage is one contributor to the glycan profiles. However, there are differences between mutants. First considering NFL, the most profound reduction in glycan processing occurred at N135, N276, N356, and N611. The latter is unusual, as the 4 gp41 glycans are usually all heavy and complex. Notably, the N637 was completely unoccupied in NFL and was only partially occupied in both the other mutants (S12B Fig). N611 was also poorly processed in I559P but not A433P, consistent with increased I559P gp41 endo H-sensitivity. This suggests that gp41 glycan sites are poorly accessible prior to gp120/gp41 processing. N463 was also unusually underprocessed in I559P. I559P’s generally poor glycan maturation may explain the increased 35O22 binding in BN-PAGE (Fig. 7C-D, lane 14). This akin to the improved 35O22 sensitivity of GnT1-modified pseudovirus, where glycan clashes are reduced [46].

### Mutant glycan profiles compared to NFL

By comparing to NFL, the effects of I559P and A433P can be understood independently (Fig. 11E-F, S1 Main glycan analysis). Both mutants increased N276 glycan processing, rescuing the glycans closer to the maturation state of the parent. N88, N160, N188, and N392 were well-differentiated in A433P. This could be a result of open trimer conformation and could account for the decreased mobility of A433P clones in SDS-PAGE (Fig. 2, lane 5). If A433P is a more “open” trimer, it makes sense that N160 is more processed as it is on gp120 monomer, where there are no constraints. N392 also appears to be regulated by compactness, as discussed above.

The most important difference is the N611 glycan which for I559P is less differentiated but is restored to a complex glycan in A433P. Data for gp41 glycans at positions 616 and 637 were not available due to proximity to N611 and/or skipping, but N625 is complex and appears unaffected in the mutants. Thus, the partial sensitivity of I559P gp41 to endo H appears to be explained by dramatically reduced glycan maturation at N611 and N637.

We observed a decrease in the fucosylated glycans in I559P compared to A433P (S12B Fig). However, all mutants showed an overall fewer fucosylated glycans than the average parent. This may simply reflect reduced glycan processing and the resulting bias towards high mannose glycans that do not exhibit these terminal modifications.

Finally, we noted that N301 glycan is an outlier in terms of its variable maturation between samples. It showed score of 10 in the 2023 sample, 17 in 2021 and was not detected in 2020 sample. Previous data suggests that this glycan is prone to “flipping”. It is often of complex type in viral Env and high mannose in soluble SOSIP [73]. Although gp160 showed the lowest score of 4 at N301, viral gp120 and gp120 monomer had intermediate scores of 7 and 8, so it is difficult to make an argument based on accessibility or else the gp120 monomer would show increased processing. The two mutants A433P and I559P also showed scores of 7, adding to this paradox. Overall, this suggests that it is subject to high variability in different preparations of the same sample, perhaps indicating a stochastic process.

## DISCUSSION

Functional Env folding involves trimerization, glycan addition and maturation, chaperone engagement, disulfide isomerization, signal peptide removal and gp120/gp41 processing. Given this complexity, a modicum of misfolded Env is perhaps inevitable. How gp160i avoids glycan maturation in the ER/Golgi is currently unknown. Revealingly, none of our mutants impacted gp160i, consistent with it being an early misfolding product that exits conventional folding before mutants can have an effect. Our BN-PAGE analysis showed that uncleaved forms of Env, including UNC, NFL, gp160m and gp160i are largely monomeric. In contrast, we recently observed that nascent gp160 extracted from cell lysates is *entirely* trimeric [28]. The reasons for this dichotomy are unclear. Possibly, trimers are stabilized by cellular factors in lysates that are largely absent by the time Env reaches the outer cell membrane. Pretreating VLPs with a crosslinker prevented some but not all UNC trimers from laterally dissociating. In contrast, cleaved trimers gained sufficient lateral stability to survive lysis.

UNC’s lateral instability may be linked to its conformational flexibility. A recent study used crosslinkers to characterize two asymmetric UNC isoforms [27]. Indeed, expression of gp160 in the presence of BMS-806, an entry inhibitor that binds the Phe43 cavity and promotes a “state 1” conformation, reduced gp120/gp41 processing and glycan maturation [27]. Here, I559P, DS and NT1-5 mutants also reduced or eliminated gp120/gp41 processing. In the case of I559P and NT1-5, BN-PAGE showed an increase in trimer, coupled with reduced monomer in the case of NT1-5. The poor gp120/gp41 processing of these mutants suggest that reduced UNC flexibility improves lateral trimer stability. I559P also improved Env expression, as appears to be common for proline mutants [74].

A key question is whether viral gp120 and gp160 glycan differences occur before or after gp120/gp41 processing. Envs that are either fully gp120/gp41 processed or completely unprocessed might help to address this point. NFL mutant is instructive. At position 356, NFL gp160 glycans were more mixed than the average parent or viral gp120 (see average sheet in S1 Main glycan analysis). At N392, NFL was largely oligomannose, unlike parent gp160, but more consistent with the average parent. This suggests the presence of a native-like conformation in NFL that would have been gp120/gp41 processed had the cleavage site had been intact. Moreover, it suggests that differences in glycan maturation precede gp120/gp41 processing. Further, this indicates at least two gp160 populations in NFL, one that is folded and dominant and others that are misfolded and acquire somewhat different glycan processing. Said another way, this implies that protein folding, not gp120/gp41 cleavage dictates glycan maturation. This does not exclude possible late maturation of some glycans, most notably of gp41 in which high mass complex glycans appear to develop only after gp120/gp41 cleavage.

The glycan analysis revealed that native Env trimers (i.e., viral gp120) exhibited more processed glycans compared to the mutants and lacked sequon skipping. In contrast, the mutants all showed large changes in gp41 glycans. They also showed increased skipping. This raises the possibility that they would induce Abs to non-encoded glycan holes if they were used as immunogens. This raises a possible scenario in which glycans progressively mature as gp160 disulfides isomerize and exchange in until they reach the canonical disulfide conformation, at which point glycans become most differentiated. A less mature glycan profile may therefore be aborted attempts to reach this final conformation and is thus a marker for misfolding. However, this is offset in globally sensitive mutants that exhibit increase processing at certain sites.

### Which modifications do we recommend for membrane Env vaccine immunogens?

For antigenic comparisons, we used virus capture, a convenient alternative to cell ELISA or flow cytometry [10,38,53]. Combining this with BN-PAGE shifts allowed us to discriminate MAb binding to trimers versus non-functional Env. We consider the antigenicity of each modification below.

#### Gp41 N-helix mutations

Given its adverse effect on gp120/gp41 processing, it is perhaps no coincidence that I559P is sometimes used in conjunction with gp120-gp41 linkers, ostensibly to achieve a “native like” conformation. The antigenic profile of I559P and NT1-5 showed some differences from parent trimers, most clearly for V2 and interface bNAbs, consistent with other studies [33,38–40]. Although some V3 non-NAb epitopes are partially exposed on soluble SOSIP trimers [33], we did not see differences in membrane trimers: V3 MAbs did not bind to either parent or I559P trimers in BN-PAGE, but VLPs were captured equivalently by V3 MAbs, presumably via non-functional Env. Lastly, UFO significantly reduced membrane trimer expression. A recent study showed that incorporating both gp120-gp41 linker and UFO into a Clade C soluble gp140 resulted in low trimer expression and reduced V2 NAb binding [75]. The same mutations, when applied to another soluble Env, resulted in a higher trimer expression [10], suggesting that UFO and/or NFL mutations are not universally adaptable.

The modest but significant 2.2-fold reduced HIV+ plasma binding to I559P (Fig. 10), suggests globally diminished Ab reactivity across multiple epitopes. Extrapolating this in a vaccine setting, I559P trimers are likely to induce some I559P-specific Abs that inefficiently cross-react with native trimers, thereby blunting potential NAb titers and reducing vaccine efficacy.

The most notable feature of I559P’s glycan profile was its greatly decreased glycan processing at N637 and to a lesser extent at N611. This may be due to the self-association of gp41 C-helices that results in reduced accessibility for glycan processing enzymes. Furthermore, glycan skipping was observed to varying degrees at N637 in NFL+ mutants, whereas all gp41 glycans are complex in the parent. The mutations create non-encoded glycan holes, like that reported in BG505 gp140 SOSIP trimer where low glycan occupancy was observed at position 611 [76]. Immunogens with such glycan holes trigger autologous Ab responses that are unable to cross-react with membrane Envs that do not have glycan holes [77]. Collectively, we advise against incorporating gp41 N-helix mutants I559P, NT1-5 and UFO for membrane trimers.

We note that some mutations directly ablate epitopes rather than mediate conformational improvement. For example, UFO may directly ablate gp41 cluster I or II non-NAb epitopes, rather than eliminate them conformationally. Hence, the lack of binding by these MAbs does not imply the Env adopts a more native conformation. Similarly, given its proximity to the 17b epitope, A433P may directly knock out 17b binding rather than prevent CD4-induced conformational change.

#### DS

The DS mutant did not stabilize the native-like membrane trimers and lacking sCD4 binding constraint suggests that the DS disulfide may not form as intended [36]. One explanation is that, like I559P, NT1-5 and UFO, DS diminishes UNC’s flexibility that is crucial for attaining correct disulfide bonding and gp120/gp41 processing [10,52]. The DS mutant was designed based on soluble SOSIP structure that conformationally differs from membrane trimers, such that positions 201 and 433 may not be proximal [31], and that these cysteines may instead react with others to form non-canonical disulfides, possibly accounting for dimers.

#### A433P

A433P caused a globally sensitive phenotype like A328G, eliminating PGT145 binding and increasing V3 MAb and CD4bs non-NAb binding. Its “open trimer” conformation led to increased glycan maturation at several variable positions, namely N88, N188, N276, N392, and N611. However, like I559P and NFL, position 637 was often skipped and/or confined to oligomannose, contrasting sharply with the parent. Therefore, N637 is commonly impacted by mutations, perhaps as its proximate position to the viral membrane can dramatically impact accessibility to glycan processing enzymes.

Globally sensitive mutants like A433P and A328G are ubiquitous among membrane Env point mutants [40,78,79] and appear to reflect misfolding. These mutants show reduced trimer in BN-PAGE, suggesting that globally sensitive trimers are laterally labile. Although I559P “rescued” A433P trimers, dimers also became prominent, suggesting misfolding. Globally sensitive mutants should be avoided in vaccine designs by monitoring for V3 MAb sensitivity [22].

#### SOS and ΔCT

Gp120 shedding appears to occur largely as a by-product of trimer synthesis, at least for primary isolates. We suggest that a fraction of misfolded Env manages to be processed into gp120/gp41, but due to misfolding, lacks sufficient cognate gp120-gp41 non-covalent associations so that gp120 immediately dissociates [80]. This leads to the exposure of gp41 stumps which are highly immunogenic and likely account for the greater HIV+ plasma reactivity with WT over SOS membrane Env (Fig. 10). To try to force a greater focus in vaccine Ab responses to NAb epitopes, we have used the SOS mutant in multiple vaccine studies [25,26,81]. SOS appears to improve trimer expression, which may simply be due to the retained gp120. Overall, the mild antigenic cost of SOS may be less important than eliminating immunodominant gp41 epitopes.

Our previous vaccine studies used ΔCT clones due to increased gp120/gp41 processing efficiency and expression. In the context of JR-FL, neutralization profiles are largely unchanged by either SOS or ΔCT. In other studies, ΔCT also did not affect NAb sensitivity [33,36,38]. However, in some contexts, these mutations cause unwanted increases NAb sensitivity, i.e., misfolding, as we found for PC64 (S6 Fig) [82]. While the full gp41 tail may be linked to a compact trimer apex, as evidenced by enhanced PG16 capture (Fig. 9), an important caveat of FL for vaccine use is its extremely poor gp120/gp41 cleavage.

We were intrigued that FL gp160 was not *detectably* bound by MAbs in BN-PAGE (Fig. 8C). This must be reconciled with the fact that the *same* MAbs can capture FL VLPs. Other BN-PAGE shifts reveal a similarly puzzling lack of MAb binding to the *monomer* (Fig. 7). We suggest that lack of MAb binding to trimers and monomers are linked. It may be no coincidence that FL is very poorly cleaved and expresses unusually high levels of gp160i compared to ΔCT clones (S4 Fig). Gp160i partly accounts for monomers in BN-PAGE. Given the predominance of gp160i over gp160m and gp120/gp41 in FL clones (S4 Fig), gp160i may also significantly account for the FL trimer in BN-PAGE. We further suggest that MAb binding to gp160i on intact VLPs is prevented by gp160i complexing with host chaperones that might be linked to its trafficking to cell supernatants. These complexes are held together by low affinity interactions that, upon VLP lysis, become dissociated, thereby allowing MAbs to bind (Fig. 8). Despite this, FL VLPs exhibited traces of functional trimer and non-functional Env that presumably is not complexed to chaperones, thereby allowing virus capture and neutralization, contrasting with BN-PAGE which is dominated by gp160i.

#### Gp120-gp41 linkers

One argument for the use of gp120-gp41 linkers in vaccines is that cellular furin processing is almost invariably incomplete, leading to an heterogeneous product. Thus, cleaved gp120/gp41 trimers will be arrayed alongside the more Ab-sensitive UNC trimers.

Multivalent vaccines such as measles, mumps, rubella (MMR) demonstrate that multiple Ab lineages can develop to distinct antigens in parallel. However, it is unknown how similar but non-identical trimer isoforms might influence Ab responses to each other. Given the sensitivity of UNC to “common in repertoire” CD4bs, V3 and other non-NAbs, a bias to UNC-reactive Abs seems almost inevitable. By analogy, glycan holes are preferential Ab targets and may delay NAb development [83,84]. For similar reasons, gp160-expressing mRNA vaccines have so far been only moderately successful [52], perhaps because non-functional Env and/or mutations create antigenic differences such that Abs insufficiently cross-react with native trimers.

Our long-standing goal is to develop a vaccine that expresses *fully* cleaved trimers - unfettered by UNC [36,39,64]. Such immunogen has yet to be tested. Overall, our data suggests that membrane Env mutations should be kept to a minimum, as any perceived gains are often countered by losses.

## MATERIALS AND METHODS

### Plasmids

i. **Env.** Abbreviated Env strain names are given first, with full names and GenBank references in parentheses: JR-FL (JR-FL, AY669728.1), PC64 (PC64 MRCA, ASP69831.1). These were expressed in plasmids pCDNA3.1 (PC64 MRCA) or pCAGGS (JR-FL). All gene synthesis, cloning and mutagenesis was performed by GenScript (USA).
ii. **Gag and Rev.** A plasmid expressing murine leukemia virus (MLV) Gag was used to produce VLPs [90]. For Env plasmids using native codons, we co-transfected pMV-Rev 0932 that expresses codon-optimized HIV-1 Rev.
iii. **NL4-3.Luc.R-E.** This plasmid is based on HIV-1 proviral clone NL4-3 in which the firefly luciferase gene replaces Nef and does not express Env or Vpr due to frameshift mutations. This plasmid is co-transfected with Env plasmids to produce pseudovirions (PVs) for neutralization assays.
iv. **VSV-G.** A plasmid expressing vesicular stomatitis virus G protein described previously [91] is used as a readout of capture virus capture assays.
v. **pQC-Fluc and pMLV GagPol.** These two plasmids express luciferase and GagPol for use in the alternative pQC-Fluc neutralization assay format.

### MAbs and soluble CD4

MAbs were obtained from their producers or the NIH AIDS Reagent Repository. These included PGT145, PG16, CH01, VRC38.01 and PC64 35K, directed to the V2 apex; 14E, 39F, 447-52D and CO11, directed to the V3 loop; 2G12 and PGT121, directed to V3-glycan epitopes; VRC01, VRC03, b12 and 15e, directed to the CD4 binding site (CD4bs); 17b, directed to a CD4-induced epitope/bridging sheet; PGT151, 35O22 and VRC34, directed to the gp120-gp41 interface [92]; 2F5, 4E10, 10e8, 7B2 and 2.2B directed to gp41 and CR3022 directed to SARS-CoV S. Four-domain soluble CD4 (sCD4) was provided by Progenics Pharmaceuticals, Inc.

### HIV+ donor plasmas

A collection of HIV-1+ plasmas we and others previously described [70] were obtained from various subtype B- and C-infected donors. Four subtype B plasmas, from United States donors 1648, 1652, 1686, 1702 and an uninfected donor (210) were obtained from Zeptometrix (Buffalo, NY). N308, a long-term nonprogressor B plasma was described previously [93]. Subtype C plasmas were purchased from the South African Blood Bank (Johannesburg). PC64 plasma was from a subtype A infected donor, provided by Elise Landais [68].

### VLP production

For VLP production, Env plasmids were co-transfected in human embryonic kidney 293T cells using polyethyleneimine (PEI Max, Polysciences, Inc.), along with the MuLV Gag plasmid and pMV-Rev 0932 for Envs carrying native codons. 48h later, supernatants were collected, precleared, filtered, and pelleted at 50,000g. Pellets were washed with PBS, recentrifuged in a microcentrifuge at 15,000rpm, and resuspended at 1,000x the original concentration in PBS.

### Endoglycosidase H (Endo H) and enterokinase digests

For endo H digests, VLPs were mixed with 2% SDS and 2-mercaptoethanol and boiled for 5 minutes. Samples were then split in half. To each sample, 2μl of endo H (1000U) (New England Biolabs) or 2μl of PBS (mock) were added and samples were incubated at 37°C for 15 minutes. Samples were then lysed and processed for SDS-PAGE-Western blot.

For enterokinase (EK) digests, VLPs were mixed with rEK 10x cleavage buffer and were digested with 1μl of enterokinase (equivalent to 2.4U; Novagen) at 37°C for 30h. Samples were then lysed and processed for reducing SDS-PAGE-Western blot.

### SDS-PAGE-Western blots

VLPs were denatured by heating in 2xLaemmli buffer containing 2-mercaptoethanol (Bio-Rad) for 10 minutes at 95°C, and proteins were resolved in 4-12% Bis-Tris NuPAGE gel (ThermoFisher). Proteins were wet transferred onto a PVDF membrane and blocked in 4% skim milk/PBST. Blots were probed for 1h at room temperature with MAb cocktails in 2% skim milk/PBST, as follows (epitopes in parentheses). Anti-gp120 MAb cocktail: 2G12 (glycan), b12 (CD4bs), 39F (V3 loop), PGT121 (N332 glycan), 14E (V3 loop). Anti-gp41 MAb cocktail: 2F5 (membrane proximal ectodomain (MPER)), 4E10 (MPER), 7B2 (cluster I), 2.2B (cluster II). After washing, blots were probed with a goat anti-human IgG alkaline phosphatase (AP) conjugate (Accurate Chemicals) at 1:5,000 in 2% skim milk/PBST for 30 minutes at room temperature. Following washing, protein bands on the blots were developed with chromogenic substrate SigmaFast BCIP/NBT (Sigma).

### Blue Native (BN) PAGE-Western Blots

VLPs were solubilized in 0.12% Triton X-100 in 1mM EDTA. An equal volume of 2x sample buffer (100mM morpholinepropanesulfonic acid (MOPS), 100mM Tris-HCl, pH 7.7, 40% glycerol, and 0.1% Coomassie blue) was added. Samples were spun to remove any debris and loaded onto a 4-12% Bis-Tris NuPAGE gel and separated for 3h at 4°C at 100V. Proteins were then transferred to PVDF membrane, de-stained, and blocked in 4% skim milk in PBST. Membranes were probed with a cocktail of MAbs 39F, 2F5, b12, 4E10, 14E, and PGT121, followed by anti-human IgG AP conjugate (Accurate Chemicals) and were developed using SigmaFast BCIP/NBT. In some experiments, Env was cross-linked on VLP membrane surfaces, prior to lysis and BN-PAGE. For this purpose, the cross-linker bis(sulfosuccinimidyl) suberate (BS^3^) (Sigma) was used, as previously described [30].

In BN-PAGE band shifts, VLPs were mixed with either sCD4 or a MAb (30μg/ml) for 1h at 37°C, followed by either washing with PBS to remove excess ligand (“Wash out” method) or no washing (“Leave in” method), and subsequent processing for BN-PAGE as above. Blots were probed in two ways: first we used anti-human IgG AP conjugate and developed the blot with substrate to check for any MAb bound to VLP Env. Next, we probed the same blot with MAb cocktail, as above, to detect Env on the blot, followed by anti-human IgG AP conjugate. Band densities were determined using ImageJ software v. 1.33u (NIH freeware; http://rsb.info.nih.gov/ij/).

### Neutralization assays

i. **NL-Luc assay.** Pseudoviruses (PV) were produced by co-transfecting 293T cells with pNL4-3.Luc.R-E and an Env plasmid using PEI Max. Briefly, PV was incubated with graded dilutions of MAbs for 1h at 37°C, then added to CF2Th.CD4.CCR5 cells, plates were spinoculated, and incubated at 37°C [46]. For SOS PV, following a 2h incubation, 5mM DTT was added for 15 minutes to activate infection. All MAb/PV mixture was replaced by fresh media (i.e., a “washout“ protocol), cultured for 3 days, and luciferase activity was measured using Luciferase Assay System (Promega).
ii. **pQC-Fluc assay.** PV were produced by co-transfecting Env plasmids with pMLV GagPol and pQC-Fluc-dIRES (abbreviated as pQC-Fluc) [22]. The resulting PV were used in neutralization assays with CF2Th.CD4.CCR5, as above.

### Virus capture assays

The antigenic properties of Env on VLP surfaces was analyzed using virus capture assay [30,94,95]. Briefly, MAbs were coated on Immulon II ELISA plates (Corning, USA) overnight at 5μg/ml in PBS. Wells were washed with PBS and blocked with 3% bovine serum albumin (Sigma) in PBS. VLPs were then added to the plate and incubated for 3h, after which the wells were washed three times with PBS. Since our Env mutants were generally non-infectious, VLPs were made to co-express VSV-G. 293T cells were added to wells to measure captured VLPs that infect via amphotropic VSV-G. Infection was detected by luciferase assay, as for neutralization.

### VLP ELISA

ELISAs were performed as described previously [96]. Briefly, Immulon II ELISA plates were coated overnight at 4°C with VLPs at 20x their concentration in transfection supernatants. Following PBS washing and blocking with 4% BSA/PBS supplemented with 10% FBS at room temperature, MAbs and HIV+ human plasmas were titrated against VLPs in 2% BSA/PBS/10% FBS. Goat anti-human IgG AP conjugate and PNPP Substrate tablets (ThermoFisher) were used to detect binding. Plates were read at 405nm. MAb concentration and plasma dilution resulting in an optical density (OD) of 0.5 (approximately 5 times above background) was recorded as its binding titer.

### Env reduction, alkylation and digestion for mass spectrometry

The following VLP samples were subjected to glycopeptide analysis: JR-FL gp160ΔCT SOS, SOS NFL, SOS NFL I559P and SOS NFL A433P. VLPs were denatured by heating in 4xLaemmli buffer containing 2-mercaptoethanol (Bio-rad) for 10 minutes at 95°C, then loaded onto 4-12% Bis-Tris NuPAGE gel. Western blot was carried out to determine the positions of gp160 (includes gp160m and gp160i), gp120 and gp41 on the gel and was used as “marker” to slice Envs from the gel. Gel slices containing either gp160, and/or gp120 and gp41 were incubated in 500 µL acetonitrile (ACN) until opaque. ACN was then removed and replaced with 50 µL 10 mM Dithiothrietol (DTT) in 100 mM ammonium bicarbonate (AmBic). Gel slices were incubated in DTT solution for 30 minutes at 56°C. Gel pieces were cooled down and washed with 500 µL ACN, then incubated in the dark for 20 minutes in 50 µL 55 mM iodoacetamide (IAA) in 100 mM AmBic. Bands were then washed once with 500 µL 100 mM AmBic and ACN. Washed bands were then incubated on ice with either α-lytic protease, trypsin or chymotrypsin (13 ng/µL) in a solution of 10 mM AmBic 10% ACN. Further enzymatic solution was added and incubated on ice for another 90 minutes, ensuring the bands were fully submerged. 10-20 µL 10 mM AmBic was added, and the mixture was then incubated overnight at 37°C. Following digestion, the gel pieces were spun down and supernatants were set aside in a fresh tube. The individual supernatants were dried using a heated vacuum centrifuge set to 30°C and resuspended in 200 μL 0.1% trifluoroacetic acid (TFA). The Oasis PRiME HLB 96-well μElution plate (Waters) was placed on a vacuum manifold set to 5” Hg. The HLB μElution wells were conditioned using 200 μL ACN, then equilibrated with 200 µL 0.1% TFA. The vacuum was turned off and the resuspended peptides were loaded into the conditioned wells. Vacuum was applied, starting at the lowest setting, and slowly increased to 5” Hg. Wells were washed with 800 μL TFA, followed by 200 μL H_2_O to remove excess salts. The collection tray was placed in the vacuum manifold and peptides were eluted using 80% ACN in 0.1% formic acid. The eluted peptides were dried prior to LC-MS analysis using a heated vacuum centrifuge set to 30°C.

### Liquid chromatography-mass spectrometry (LC-MS) glycopeptide analysis

Peptides were dried then resuspended in 0.1% formic acid and analyzed by nanoLC-ESI MS with an Ultimate 3000 HPLC (Thermo Fisher Scientific) system coupled to an Orbitrap Eclipse mass spectrometer (Thermo Fisher Scientific) using stepped higher energy collision-induced dissociation (HCD) fragmentation. Peptides were separated using an EasySpray PepMap RSLC C18 column (75µm × 75cm). A trapping column (PepMap 100 C18 3μM 75μM × 2cm) was used in line with the LC prior to separation with the analytical column. LC conditions were as follows: 280 minute linear gradient consisting of 4-32% ACN in 0.1% formic acid over 260 minutes, followed by 20 minutes of alternating 76% ACN in 0.1% formic acid and 4% ACN in 0.1% formic acid to ensure all the sample elutes from the column. The flow rate was set to 300nL/min. The spray voltage was set to 2.7 kV and the temperature of the heated capillary was set to 40°C. The ion transfer tube temperature was set to 275°C. The scan range was 375−1500 m/z. Stepped HCD collision energy was set to 15%, 25% and 45% and the MS2 for each energy was combined. Precursor and fragment detection were performed with an Orbitrap at a resolution MS1=120,000, MS2=30,000. The AGC target for MS1 was set to standard and injection time set to auto which involves the system setting the two parameters to maximize sensitivity while maintaining cycle time.

### Site-specific glycan classification

Glycopeptide fragmentation data were extracted from the raw file using Byos (Version 3.5; Protein Metrics Inc.). Glycopeptides were evaluated in reference to UniProtKB Q6BC19 (ectodomain of JR-FL gp160ΔCT). All samples carry mutations SOS (A501C, T605C) and E168K and N189A, and others, as denoted in the .txt files found in the MassIVE database (MSV000088108). Data were evaluated manually for each glycopeptide. A peptide was scored as true-positive when the correct b and y fragment ions were observed, along with oxonium ions corresponding to the glycan identified. The MS data was searched using the Protein Metrics “N-glycan 309 mammalian no sodium” library with sulfated glycans added manually. All charge states for a single glycopeptide were summed. The precursor mass tolerance was set at 4 ppm and 10 ppm for fragments. A 1% false discovery rate (FDR) was applied. Glycans were categorized according to the composition detected.

HexNAc(2)Hex(10+) was defined as M9Glc, HexNAc(2)Hex(9−3) was classified as M9 to M3. Any of these structures containing a fucose were categorized as FM (fucosylated mannose). HexNAc(3)Hex(5−6)X was classified as Hybrid with HexNAc(3)Hex(5-6)Fuc(1)X classified as Fhybrid. Complex glycans were classified according to the number of HexNAc subunits and the presence or absence of fucosylation. As this fragmentation method does not provide linkage information, compositional isomers are grouped. For example, a triantennary glycan contains HexNAc(5) but so does a biantennary glycans with a bisect. Core glycans refer to truncated structures smaller than M3. M9Glc-M4 were classified as oligomannose glycans. Glycans containing at least one sialic acid were categorized as NeuAc and at least one fucose residue in the “fucose” category.

Glycans were categorized into I.D.s ranging from 1 (M9Glc) to 19 (HexNAc(6+)(F)(x)). These values were multiplied by the percentage of the corresponding glycan divided by the total glycan percentage excluding unoccupied and core glycans to give a score corresponding to the most prevalent glycan at a given site. Arithmetic score changes were then calculated from the subtraction of these scores from one sample against others, as specified.

### Construction of trimer model and cognate glycans

The model representation of the JR-FL gp160ΔCT SOS E168K+N189A trimer was constructed using SWISS-MODEL based on an existing structure of the 426c DS-SOSIP D3 trimer (pdb: 6MYY). Glycans were modelled on to this structure based on the most abundant glycoform identified from site-specific glycan analysis using WinCoot version 0.9.4.1 and PyMOL version 2.5.0. For sites that were not identified, a Man9GlcNAc2 glycan was modelled. Conditional color formatting was used to illustrate the predominant glycoforms of modeled glycans, as follows: green (high mannose), white (hybrid) and magenta (complex). Glycan score difference between clones were represented with conditional color formatting as follows: red (negative scores), white (zero score) and blue (positive scores).

### Statistical analysis

All graphs were generated and analyzed using Prism (version 9.5.1).

## ACKNOWLEDGEMENTS

We thank Bill Schief for discussions and Elise Landais for providing PC64 reagents.

